# A Multidimensional Systems Biology Analysis of Cellular Senescence in Ageing and Disease

**DOI:** 10.1101/743781

**Authors:** Roberto A. Avelar, Javier Gómez Ortega, Robi Tacutu, Eleanor Tyler, Dominic Bennett, Paolo Binetti, Arie Budovsky, Kasit Chatsirisupachai, Emily Johnson, Alex Murray, Samuel Shields, Daniela Tejada-Martinez, Daniel Thornton, Vadim E. Fraifeld, Cleo L. Bishop, João Pedro de Magalhães

**Affiliations:** Integrative Genomics of Ageing Group, Institute of Ageing and Chronic Disease, University of Liverpool, Liverpool L7 8TX, UK; Centre for Cell Biology and Cutaneous Research, Blizard Institute, Barts and The London School of Medicine and Dentistry, Queen Mary University of London, London E1 2AT, UK; Research and Development Authority, Barzilai Medical Center, Ashkelon, Israel; The Shraga Segal Department of Microbiology, Immunology and Genetics, Faculty of Health Sciences, Center for Multidisciplinary Research on Aging, Ben-Gurion University of the Negev, Beer Sheva 8410501, Israel; School of Biological Sciences, Monash University, Melbourne, Victoria 3800, Australia; Computational Biology of Aging Group, Institute of Biochemistry, Romanian Academy, Bucharest, 060031, Romania; Chronos Biosystems SRL, Bucharest, 060117, Romania; Instituto de Ciencias Ambientales y Evolutivas, Facultad de Ciencias, Universidad Austral de Chile, Independencia 631, Valdivia, Chile

## Abstract

Cellular senescence, a permanent state of replicative arrest in otherwise proliferating cells, is a hallmark of ageing and has been linked to ageing-related diseases like cancer. Senescent cells have been shown to accumulate in tissues of aged organisms which in turn can lead to chronic inflammation. Many genes have been associated with cell senescence, yet a comprehensive understanding of cell senescence pathways is still lacking. To this end, we created CellAge (http://genomics.senescence.info/cells), a manually curated database of 279 human genes associated with cellular senescence, and performed various integrative and functional analyses. We observed that genes promoting cell senescence tend to be overexpressed with age in human tissues and are also significantly overrepresented in anti-longevity and tumour-suppressor gene databases. By contrast, genes inhibiting cell senescence overlapped with pro-longevity genes and oncogenes. Furthermore, an evolutionary analysis revealed a strong conservation of senescence-associated genes in mammals, but not in invertebrates. Using the CellAge genes as seed nodes, we also built protein-protein interaction and co-expression networks. Clusters in the networks were enriched for cell cycle and immunological processes. Network topological parameters also revealed novel potential senescence-associated regulators. We then used siRNAs and observed that of 26 candidates tested, 19 induced markers of senescence. Overall, our work provides a new resource for researchers to study cell senescence and our systems biology analyses provide new insights and novel genes regarding cell senescence.

## INTRODUCTION

In the 1960s, Leonard Hayflick and Paul Moorhead demonstrated that human fibroblasts reached a stable proliferative growth arrest between their fortieth and sixtieth divisions (Hayflick and Moorhead, 1961). Such cells would enter an altered state of “replicative senescence,” subsisting in a non-proliferating, metabolically-active phase with a distinct vacuolated morphology (Kuilman et al., 2010). This intrinsic form of senescence is driven by gradual replicative telomere erosion, eventually exposing an uncapped free double-stranded chromosome end and triggering a permanent DNA damage response (DDR) (d’Adda di Fagagna et al., 2003; Herbig et al., 2004). Additionally, acute premature senescence can occur as an antagonistic consequence of genomic, epigenomic, or proteomic damage, driven by oncogenic factors, oxidative stress, or radiation (de Magalhaes and Passos, 2018). Initially assumed to be mainly an evolutionary response to reduce mutation accrual and subsequent tumorigenesis, the pleiotropic nature of senescence has since also been positively implicated in embryogenesis (Munoz-Espin et al., 2013; Storer et al., 2013), wound healing (Demaria et al., 2014) and immune clearance (Burton and Stolzing, 2018; Kang et al., 2011). By contrast, the gradual accumulation and chronic persistence of senescent cells with time promotes deleterious effects that are considered to accelerate deterioration and hyperplasia in ageing (Campisi, 2013). Senescent cells secrete a cocktail of inflammatory and stromal regulators – denoted as the senescence-associated secretory phenotype, or SASP – which adversely impact neighbouring cells, the surrounding extracellular matrix, and other structural components, resulting in chronic inflammation, the induction of senescence in healthy cells, and vulnerable tissue (Acosta et al., 2013; van Deursen, 2014). Transgenic mice expressing transgene INK-ATTAC, which induces apoptosis of senescence cells, also increases lifespan and improves healthspan (Baker et al., 2016). It is, therefore, no surprise that in recent years gerontology has heavily focused on the prevention or removal of senescent cells as a means to slow or stop ageing and related pathologies (Baar et al., 2017; Baker et al., 2011; Yosef et al., 2016).

Research has sought to ascertain the genetic programme and prodrome underlying the development and phenotype of senescent cells (Vaziri and Benchimol, 1998). Expedited by recent advances in genomic and transcriptomic sequencing, alongside high throughput genetic screens, a wealth of publicly available data now exists which has furthered the understanding of senescence regulation (Hernandez-Segura et al., 2017; Lafferty-Whyte et al., 2010). Unfortunately, despite our increasing knowledge of CS, determining whether a cell has senesced is not clear-cut. Common senescence markers used to identify CS *in vitro* and *in vivo* include senescence-associated β-galactosidase (SA-β-gal) and cyclin-dependent kinase inhibitor 2A (p16^INK4A^) (Chandler and Peters, 2013; Dimri et al., 1995; Sharpless and Sherr, 2015). However, β-galactosidase activity has been detected in other cell types too such as macrophages, osteoclasts, and cells undergoing autophagy (Bursuker et al., 1982; Kopp et al., 2007; Young and Narita, 2010). Furthermore, some forms of senescence are not associated with p16^INK4A^ expression, whilst there can also be p16^INK4A^ in non-senescent cells (Herbig et al., 2004; Witkiewicz et al., 2011). As such, there are now over two hundred genes implicated in cellular senescence in humans alone. Therefore, it is necessary to conglomerate this data into a purposefully designed database.

Gene databases are highly useful for genomic computational analyses, as exemplified by the Human Ageing Genomic Resources (HAGR) (Tacutu et al., 2018). HAGR provides a portal where users can study ageing from various perspectives using structured data such as longevity-modulating genetic interventions, age-related molecular changes, longevity-associated gene variants, organism data, and drugs that increase lifespan. CellAge builds on these HAGR facilities to provide a means of studying cell senescence in the context of ageing or as a standalone resource; the expectation is that CellAge will now provide the basis for processing the discrete complexities of cellular senescence on a systematic scale.

Our recent understanding of biological networks has led to new fields, like network medicine (Barabasi et al., 2011). Biological networks can be built using protein interaction and gene co-expression data. A previous paper used protein-protein interactions to build genetic networks identifying potential longevity genes along with links between genes and ageing-related diseases (Budovsky et al., 2007). Here, we present the network of proteins and genes co-expressed with the CellAge senescence genes. Assaying the networks, we find links between senescence and immune system functions, and find genes highly connected to CellAge genes under the assumption that a guilt-by-association approach will reveal genes with similar functions (Vidal et al., 2011).

In this study, we look at the broad context of Cellular Senescence (CS) genes – their association with ageing and ageing-related diseases, functional enrichment, evolutionary conservation, and topological parameters within biological networks – to further our understanding of the impact of CS in ageing and diseases. Using our networks, we generate a list of 30 potential novel CS regulators and experimentally validate 26 genes using siRNAs, identifying 19 potential senescence inhibitors.

## RESULTS

### The CellAge Database

The CellAge website can be accessed at http://genomics.senescence.info/cells/. Figure 1A presents the main CellAge data browser, which allows users to surf through the available data. The browser includes several columns with information that can be searched and filtered efficiently. Users can search for a comma-separated gene list or for individual genes. Once selected, a gene entry page with more detailed description of the experimental context will open.

**Figure 1.**
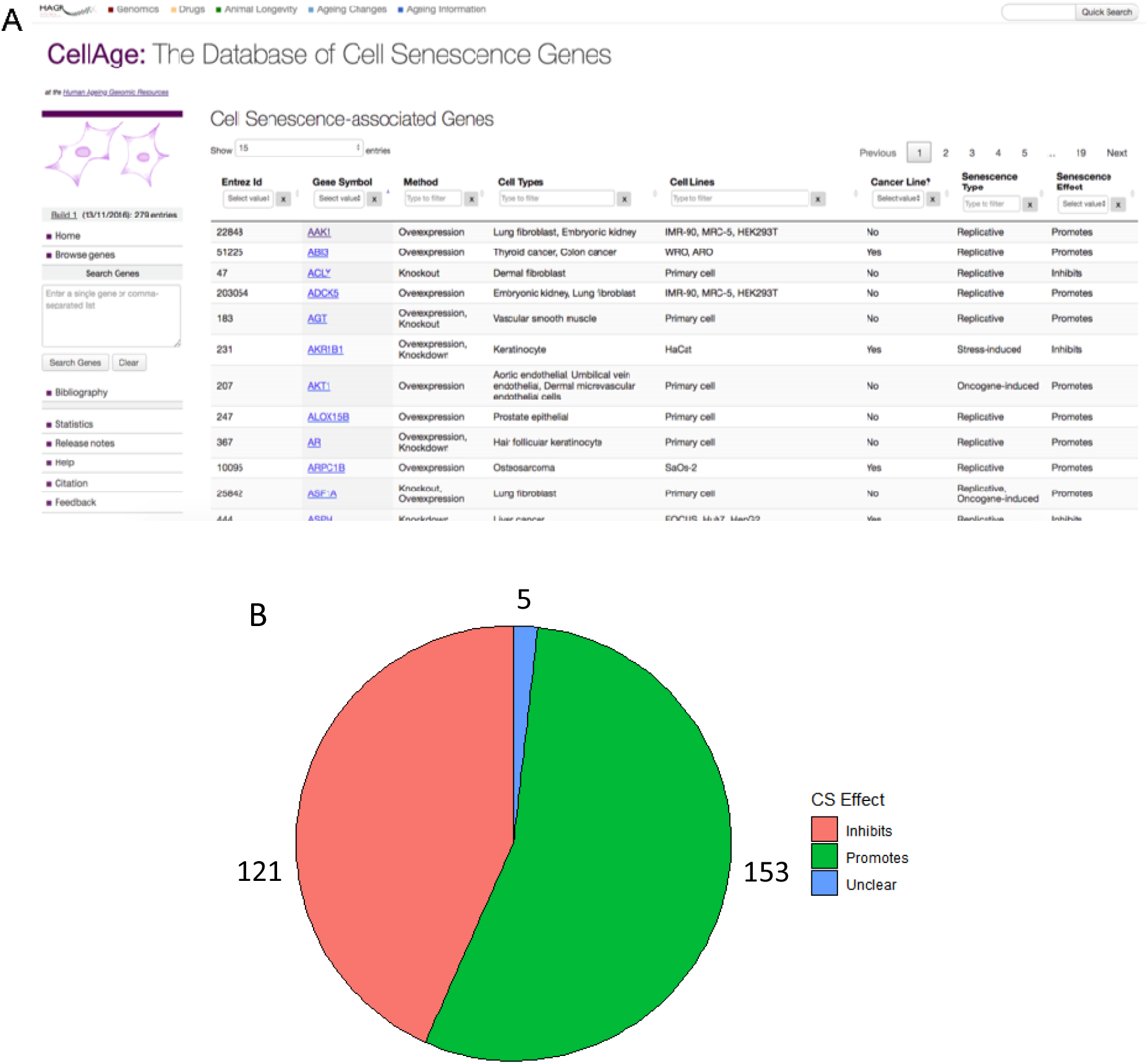
**(A) The CellAge Database of CS genes.** The main data browser provides functionality to filter by multiple parameters like cell line and senescence type and select genes to view details and links with other ageing-related genes on the HAGR website **(B) Breakdown of the effects all 279 CellAge genes have on CS.** Genes marked as ‘unclear’ both promote and inhibit senescence depending on biological context.

CellAge was compiled following a scientific literature search of gene manipulation experiments in primary, immortalized, or cancer human cell lines that caused cells to promote or inhibit CS. The first CellAge build comprises 279 distinct CS genes, of which 232 genes affect replicative CS, 34 genes are associated with stress-induced CS, and 28 genes are associated with oncogene-induced CS. Of the 279 total genes, 153 genes promote CS, 121 inhibit it and five genes have unclear effects, both promoting and inhibiting CS depending on experimental conditions (Figure 1B). The genes in the dataset are also classified according to the experimental context used to determine these associations. We have also performed a meta-analysis to identify the genetic signatures of replicative CS, and found 526 overexpressed and 734 underexpressed genes (Chatsirisupachai et al., 2019). These gene signatures are also available on the CellAge website. Of the 279 CellAge genes, 44 genes were present in the signatures of CS (15.8%). This overlap was significant (P-value = 1.62e-08; Fisher’s exact test). While 13 of the CellAge promoters of CS significantly overlapped with the overexpressed signatures of CS (8.5%, P=2.06e-06, Fisher’s exact test), only 7 overlapped with the underexpressed signatures (4.6%, P=5.13e-01, Fisher’s exact test). The CellAge inhibitors of CS significantly overlapped with both the overexpressed signatures of CS (n=7, 5.8%, P=4.08e-02, Fisher’s exact test) and underexpressed signatures of CS (n=17, 14%, P=2.06e-06, Fisher’s exact test).

### CellAge Gene Functions

High quality curated datasets enable systematic computational analyses (Barardo et al., 2017; Fernandes et al., 2016). Since we are interested in learning more about the underlying processes and functionality shared by human CS genes, we started by exploring the functional enrichment within the CellAge dataset.

Using the database for annotation, visualisation and integrated discovery – DAVID Version 6.8 (Huang da et al., 2009a, b), we found that genes in CellAge are enriched with several clusters associated with Protein Kinase Activity, Transcription Regulation, DNA-binding, DNA damage repair and Cell cycle regulation in cancer. In particular, genes that promote senescence were more associated with promoting transcription, while genes that inhibit senescence were more associated with repressing transcription. Furthermore, we found that promoters of senescence were significantly associated with VEGF and TNF signalling pathways (p<0.01, Fisher’s exact test with Benjamini-Hochberg correction) (SI Table 1; SI Table 2).

### Evolutionary Conservation of CellAge Genes in Model Organisms

Next, we looked at the conservation of CellAge genes across a number of mammalian and non-mammalian model organisms with orthologues to human CellAge genes using Ensembl BioMart (Version 96) (Smedley et al., 2015) in order to understand the genetic conservation of CS processes. There was a significantly higher number of human orthologues for CellAge genes than for other protein-coding genes in mouse, rat, and monkey, while non-mammalian species did not show significant conservation of CellAge genes (two-tailed z-test with BH correction) (SI Figure 1A; SI Table 3). Interestingly, previous studies have found that longevity-associated genes (LAGs) are substantially overrepresented from bacteria to mammals, and that the effect of LAG overexpression in different model organisms was mostly the same (Yanai et al., 2017). It remains unclear what the evolutionary origin of most of the CellAge genes is or why they are not present in more evolutionarily distant organisms. Unique evolutionary pressures could have played an important role in the evolution of CellAge genes in mammals. However, somatic cells in *C. elegans* and *Drosophila* are post mitotic and lack an equivalent CS process, which could explain why the CellAge genes are not conserved. We further compared the conservation of CellAge promoters and inhibitors of CS and found that while the promoters were significantly conserved in the mammal model organisms, the inhibitors were not (SI Figure 1B).

We also report the number of orthologous CellAge genes present in 24 mammal species using the OMA standalone software v. 2.3.1 algorithm (Altenhoff et al., 2015) (SI Figure 1C). From 279 CellAge genes, we report 271 orthogroups (OG) (SI Cellage_Orthologous-Groups). 22 OGs were conserved in the 24 mammals, including the following genes: *DEK*, *BRD7*, *NEK4*, *POT1*, *SGK1*, *TLR3*, *CHEK1*, *CIP2A*, *EWSR1*, *HDAC1*, *HMGB1*, *KDM4A*, *KDM5B*, *LATS1*, *MORC3*, *NR2E1*, *PTTG1*, *RAD21*, *NFE2L2*, *PDCD10*, *PIK3C2A*, and *SLC16A7* (SI Table 4). Within the long-lived mammal genomes we analysed (human, elephant, naked mole rat, bowhead whale, and little brown bat), we found 128 OG CellAge genes (SI CellAge_Orthologous-Groups; genomes available in SI Table 5). However, finding OGs is dependent on genome quality and annotations, and higher quality genomes would likely yield more OGs.

### CellAge vs Human Orthologues of Longevity-Associated Model Organism Genes

To understand how senescence is linked to genetics of ageing processes, we looked at the intersection of CellAge genes and the 869 genes in the human orthologues of model organisms’ longevity-associated genes (LAGs) dataset, collected based on quantitative changes in lifespan (Fernandes et al., 2016). Like CellAge, where genes are classified based on whether their upregulation promotes, inhibits, or has an unknown impact on CS, the longevity orthologues dataset also provides information on the effect of upregulation of its genes, namely whether it promotes (pro, 421) or inhibits (anti, 448) longevity (SI Table 6; SI Figure 2).

The CS promoters statistically overlapped with the anti-longevity genes and not with the pro-longevity genes (anti: n=9, ∼6%, P=7.30e-03; pro: n=6, ∼4%, P=1.02e-01, hypergeometric distribution with Bonferroni correction). We noted an inverse result with the inhibitors of CS, where there was a much greater overlap between the CellAge inhibitors and the pro-longevity genes, resulting in the smallest p-value of all the overlaps (n=18, ∼15%, P=6.95e-11, hypergeometric distribution with Bonferroni correction). However, there was also a significant overrepresentation of genes inhibiting the CS process within the anti-longevity genes (n=7, ∼6%, P= 1.84e-02, hypergeometric distribution with Bonferroni correction). It is possible that some of the pathways the CS inhibitors are associated with increase longevity, whereas other pathways have anti-longevity effects. Overall, these results highlight a statistically significant association between CS and the ageing process and suggest a potential inverse relationship between CS and longevity, at least for some pathways. Gene overlaps are available in SI Table 7.

### CellAge Genes Differentially Expressed with Age

In another work we performed a meta-analysis to find molecular signatures of ageing derived from humans, rats, and mice (Palmer et al., in preparation). To investigate how the expression of CellAge genes changes with age, we looked for CellAge genes which either promote (153) or inhibit (121) senescence within the list of ageing signatures. The genes overexpressed with age (449) had a significant overlap with the CellAge genes (CS promoters: n = 17, ∼11%, P=1.95e-07; CS inhibitors: n=9, ∼7%, P=3.29e-03, Fisher’s exact test with BH correction) while the genes underexpressed with age (162) did not (CS promoters: n=0, P=6.39e-01; CS inhibitors: n=3, ∼3%, P=1.13e-01). The overexpressed genetic signatures of replicative CS (526) also significantly overlapped with the overexpressed signatures of ageing (n=60, ∼11%, P=6.25e-25), but not the underexpressed signatures of ageing (n=3, ∼1%, P=8.03e-1). Finally, the underexpressed signatures of replicative CS (734) did not significantly overlap with the overexpressed (n=18, ∼3%, P=8.03e-1) or underexpressed (n=9, ∼1%, P=4.06e-1) signatures of ageing.

Using all protein-coding genes as a background list, we further examined the CS promoters overexpressed with age for functional enrichment using WebGestalt to determine if specific pathways were enriched (Wang et al., 2017). In parallel, we performed this analysis using the genes which overlapped between CellAge inhibitors and genes overexpressed with age. Seventy-three GO terms were significantly enriched for the overlap between CellAge senescence promoters and age upregulated genes (p<0.05 Fisher’s exact test with BH correction) (SI Table 8). After clustering GO terms with REVIGO, we found groups enriched for regulation of apoptotic processes, response to lipid, epithelium development, rhythmic process, circadian rhythm, cytokine metabolism, and cell-substrate adhesion (SI Figure 3A) (Supek et al., 2011). Seventy-one enriched GO terms for the overexpressed signatures of CS overexpressed with age were clustered using REVIGO, resulting in enriched terms relating to leukocyte activation, aging, response to beta-amyloid, and cell proliferation (SI Table 9; SI Figure 3B). No GO terms were significantly enriched for the promoters of CS underexpressed with age, the inhibitors of CS differentially expressed with age, the underexpressed signatures of CS differentially expressed with age, or the overexpressed signatures of CS underexpressed with age.

### Tissue-Specific CS Gene Expression and Differential Expression of CS Genes in Human Tissue with Age

The Genotype-Tissue Expression (GTEx) project (v7, January 2015 release) contains expression data from 53 different tissue sites collected from 714 donors ranging from 20 to 79 years of age which can be grouped into 26 tissue classes (Consortium, 2013). We asked if CellAge genes and differentially expressed signatures of CS were expressed in a tissue-specific manner (Palmer et al., manuscript in preparation), and determined how CS gene expression changes across different tissues with age (Chatsirisupachai et al., 2019).

We first examined tissue-specific CS expression, and found that CellAge genes were either expressed in a tissue-specific manner less than expected by chance, or in line with expectations; in other words, the majority of CellAge genes tended to be expressed across multiple tissues (SI Figure 4A; SI Table 10). Testis was the only tissue with significant differences between the actual and expected number of tissue-specific CellAge genes expressed (less tissue-specific genes than expected by chance, p<0.05, Fisher’s exact test with BH correction). The underexpressed signatures of CS were significantly less tissue-specific in the testis and liver, while the overexpressed signatures of CS were significantly less tissue-specific in the brain, liver, pituitary, and skin, and more tissue-specific in blood. We also compared the ratio of tissue-specific to non-tissue-specific genes in the CS datasets to all protein-coding genes. While ∼25% of all protein-coding genes are expressed in a tissue-specific manner, only ∼10% of CellAge genes and ∼11% of signatures of CS are expressed in a tissue-specific manner (SI Figure 4B), significantly less than expected by chance (p=2.52e-12 and 3.93e-48 respectively, Fisher’s exact test with BH correction).

Then, we examined the differential expression of CS genes with age in different tissues. Using a previously generated gene set of differentially expressed genes (DEGs) with age in 26 tissues on GTEx (Chatsirisupachai et al., 2019; Consortium, 2013), we found overlaps with 268 CellAge promoters and inhibitors of CS present in the gene expression data (Figure 2A). The process of finding DEGs with age filters out lowly expressed genes, which explains the 11 missing CellAge CS regulators. Overall, senescence promoters were overexpressed across different tissues with age, although none of the overlaps were significant after FDR correction (Fisher’s exact test with BH correction, p<0.05) (SI Table 11). There was the opposite trend in the inhibitors of CS, where there was noticeably less overexpression of CS inhibitors with age, although these overlaps were also not significant after FDR correction. 1,240 differentially expressed signatures of CS were also overlapped with the GTEx ageing DEGs in 26 human tissues, including 9 tissues previously analysed (Figure 2B) (Chatsirisupachai et al., 2019). The overexpressed signatures of CS were significantly overexpressed across multiple tissues with age, and only significantly underexpressed with age in the brain and uterus (p<0.05, Fisher’s exact test with BH correction) (SI Table 12). Furthermore, the underexpressed signatures of CS trended towards overexpressing less than expected by chance across multiple tissues with age, although these overlaps were only significant after FDR adjustment in the colon and nerve, while the underexpressed signatures of CS were significantly overexpressed more than expected in the uterus. Finally, the underexpressed signatures of CS were underexpressed with age more than expected by chance in the colon, lung and ovary, and underexpressed with age less than expected by chance in the brain. We also compared the ratio of differentially expressed to non-differentially expressed CS genes in at least one tissue with age to the equivalent ratio in all protein-coding genes (SI Figure 5A) (see Overlap Analysis in materials and methods). We found that ∼64% of all protein-coding genes did not significantly change expression with age in any human tissues, while ∼19% were overexpressed, and ∼17% were underexpressed (∼7% were both overexpressed and underexpressed across multiple tissues) (SI Figure 5; SI Table 13 and 14). For the CellAge genes, the number of promoters of CS significantly overexpressed with age in at least one tissue was significantly higher than the genome average (n=50, ∼30%, p=1.5e-3, Fisher’s exact test with BH correction). The promoters of CS underexpressed with age and the inhibitors of CS differentially expressed with age were not significantly different than the protein-coding average. We also compared the number of signatures of CS differentially expressed with age in at least one tissue to the protein-coding genome average. The overexpressed signatures of CS were significantly differentially expressed with age compared to all protein-coding genes, whereas the number of underexpressed signatures of CS were underexpressed with age more than expected by chance.

**Figure 2.**
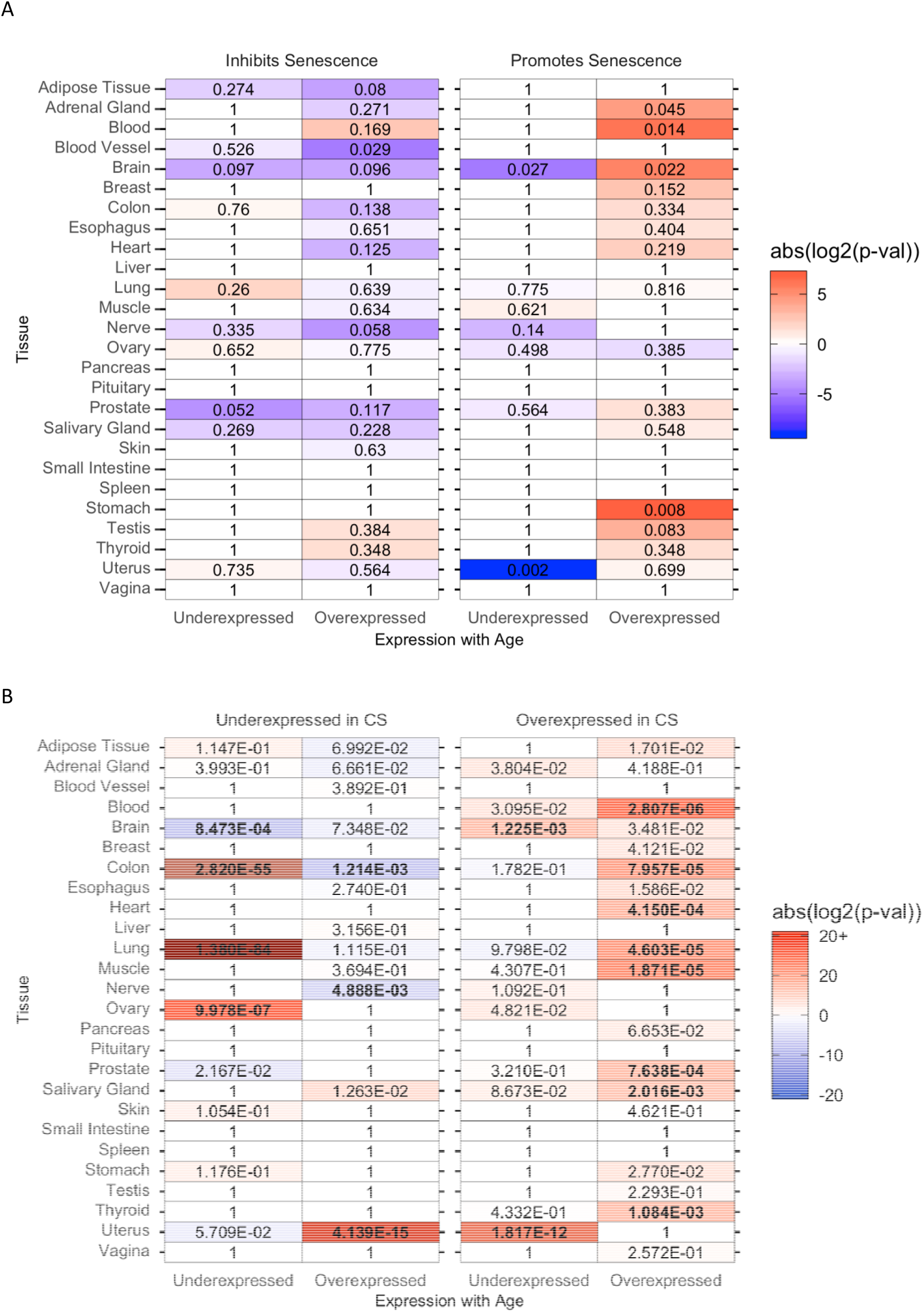
Differential expression of (A) CellAge Promoters and Inhibitors of CS and (B) differentially expressed signatures of CS in human tissue with age. Red values represent the absolute log2(p value) and indicate that there were more genes differentially expressed with age than expected by chance. Blue values indicate that there were less genes differentially expressed with age than expected by chance (negative absolute log2(p value)). Numbers indicate p-values (Fisher’s exact test), while bold p-values are significant after FDR correction (BH) (SI Table 11; SI Table 12).

The overall fold change (FC) with age of the CS genes was also compared to the FC with age of all protein-coding genes for each tissue in GTEx (Figure 3A; SI Table 15). The median log_2_FC with age of the CellAge CS promoters and the overexpressed signatures of CS was greater than the genome median for the majority of tissues on GTEx, although the difference in log_2_FC distribution with age between the promoters of CS and all protein-coding genes was only significant in seven tissues (Wilcoxon rank sum test with BH correction, p<0.05). The median log_2_FC with age of the CellAge inhibitors of CS and the underexpressed signatures of ageing was smaller than the genome median in the majority of tissues, showcasing the opposite trend to the promoters of CS and overexpressed signatures of CS. However, the only tissues with significantly different distributions of log_2_FC with age for the inhibitors of CS were the skin and oesophagus, where the median log_2_FC distribution was significantly less than the genome average, and the salivary gland, where the median log_2_FC distribution was significantly more than the genome average. We also found that the distribution of log_2_FC with age of the differentially expressed signatures of CS significantly changed in opposite directions with age in 14 tissues. Interestingly, this trend was present even in the adrenal gland and uterus, where the signatures of CS changed with age in the opposite direction to the majority of other tissues.

**Figure 3.**
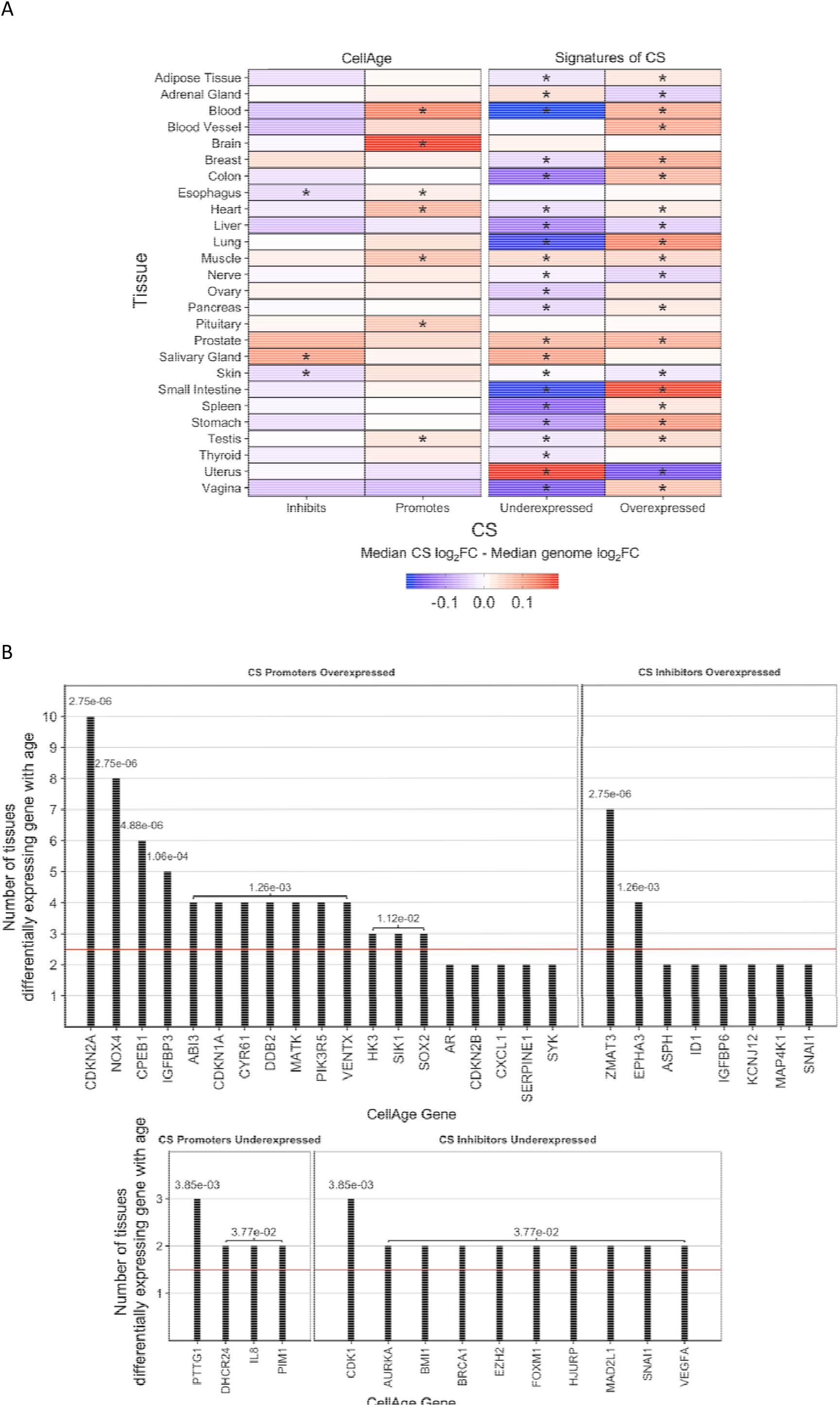
**(A) Comparison of the median log_2_FC and distribution of log_2_FC with age between the CS genes and all protein coding genes in human tissue.** Red tiles indicate that the median log_2_FC of the CS genes is higher than the median log_2_FC of all protein-coding genes for that tissue, while blue tiles indicate that the median log_2_FC of the CS genes is lower than the median genome log_2_FC. Labels indicate significant differences between the log_2_FC distribution with age of CS genes and the log_2_FC distribution with age of all protein-coding genes for that tissue (* -p<0.05, Wilcoxon rank sum test with BH correction) (SI Table 15). **(B) CellAge genes differentially expressed in multiple tissues with age.** Numbers indicate p-values while black bars above the red line indicate the CellAge gene was differentially expressed with age in more tissues than expected by chance (p<0.05, gene expression tissue overlap simulations with BH correction) (SI Table 16 – 19).

The expression of the majority of CS genes does not change with age (SI Figure 5), yet a significant number of CS genes trend towards differential expression with age across multiple tissues in humans (Figure 2). We ran 10,000 simulations on the GTEx RNA-seq data to determine the likelihood of a CS gene differentially expressing with age in more than one tissue by chance (see simulation of CS gene expression in human ageing in materials and methods). The likelihood of a CellAge gene overexpressing with age in more than three tissues and underexpressing with age in more than two tissues by chance was less than 0.05 (p<0.05, CS gene expression simulations with BH correction) (SI Table 16; Figure 3B). CS promoters overexpressed in significantly more tissues with age than expected by chance included *CDKN2A*, *NOX4*, *CPEB1*, *IGFBP3*. *ABI3*, *CDKN1A*, *CYR61*, *DDB2*, *MATK*, *PIK3R5*, *VENTX*, *HK3*, *SIK1*, and *SOX2*, while *PTTG1*, *DHCR24*, *IL8*, and *PIM1* were underexpressed in significantly more tissues (SI Table 17). *ZMAT3* and *EPHA3* were the two CS inhibitors overexpressed in significantly more tissues with age than expected by chance, while *CDK1*, *AURKA*, *BMI1*, *BRCA1*, *EZH2*, *FOXM1*, *HJURP*, *MAD2L1*, *SNAI1*, and *VEGFA* were underexpressed in significantly more tissues. We also performed simulations to determine the likelihood of a signature of CS differentially expressing with age in multiple human tissues by chance (SI Table 18). Less than 5% of the signatures of CS overexpressed with age in more than three tissues or underexpressed with age in more than two tissues. A total of 46 CS signatures (29 overexpressed signatures of CS, 17 underexpressed signatures of CS) were overexpressed with age in significantly more tissues than expected by chance, and 139 CS signatures were underexpressed in more tissues than expected by chance (26 overexpressed signatures of CS, 113 underexpressed signatures of CS) (SI Table 19).

### Do CS and Longevity Genes Associate with Ageing-Related Disease Genes?

A previous paper (Fernandes et al., 2016) grouped 769 ageing-related diseases (ARDs) into 6 NIH Medical Subject Headings (MeSH) classes (Dhammi and Kumar, 2014) based on data from the Genetic Association Database (Becker et al., 2004): cardiovascular diseases (CVD), immune system diseases (ISD), musculoskeletal diseases (MSD), nutritional and metabolic diseases (NMD), neoplastic diseases (NPD), and nervous system diseases (NSD). The same approach was used to build the HAGR ageing-related disease genes selection tool (http://genomics.senescence.info/diseases/gene_set.php), which we used to obtain the ARD genes for each disease class and overlap with the CellAge genes.

There were links between the CellAge genes and NPD genes, which is expected given the anti-tumour role senescence has (SI Table 20). Without accounting for publication bias, all ARD classes are significantly associated with CellAge genes, with lower commonalities with diseases affecting mostly non-proliferating tissue such as NSD. NPD genes are even more overrepresented in the GenAge human dataset, which could suggest commonality between ageing and senescence through cancer-related pathways. Both the strong association of NPD genes with GenAge and senescence, and the strong link between GenAge and all ARD classes is interesting. Indeed, longevity-associated genes have been linked to cancer-associated genes in previous papers (Budovsky et al., 2009). Considering ageing is the leading risk factor for ARD (Kennedy et al., 2014; Niccoli and Partridge, 2012), the results from GenAge support the previously tested conjecture that there are (i) at least a few genes shared by all or most ARD classes; and (ii) those genes are also related to ageing in general (Fernandes et al., 2016). We also looked for genes that are shared across multiple disease classes and also recorded as CS genes. CellAge genes shared across multiple ARD classes included *VZGFA* and *IFNG* (5 ARD classes), *SERPINE1*, *MMP9*, and *AR* (4 ARD classes), and *CDKN2A* (3 ARD classes). Results are summarised in SI Figure 6.

### Are CS Genes Associated with Cancer Genes?

Cellular senescence is widely thought to be an anti-cancer mechanism (de Magalhaes, 2013). Therefore, the CellAge senescence promoters and inhibitors of senescence were overlapped with oncogenes from the tumour suppressor gene (TSG) database (TSGene 2.0) (n=1,018) (Zhao et al., 2013) and the ONGene database (n=698) (Liu et al., 2017) (SI Table 21 and 22 respectively). The number of significant genes overlapping are shown in Figure 4A, while the significant p-values from the overlap analysis are shown in Figure 4B (p<0.05, Fisher’s exact test with BH correction).

**Figure 4.**
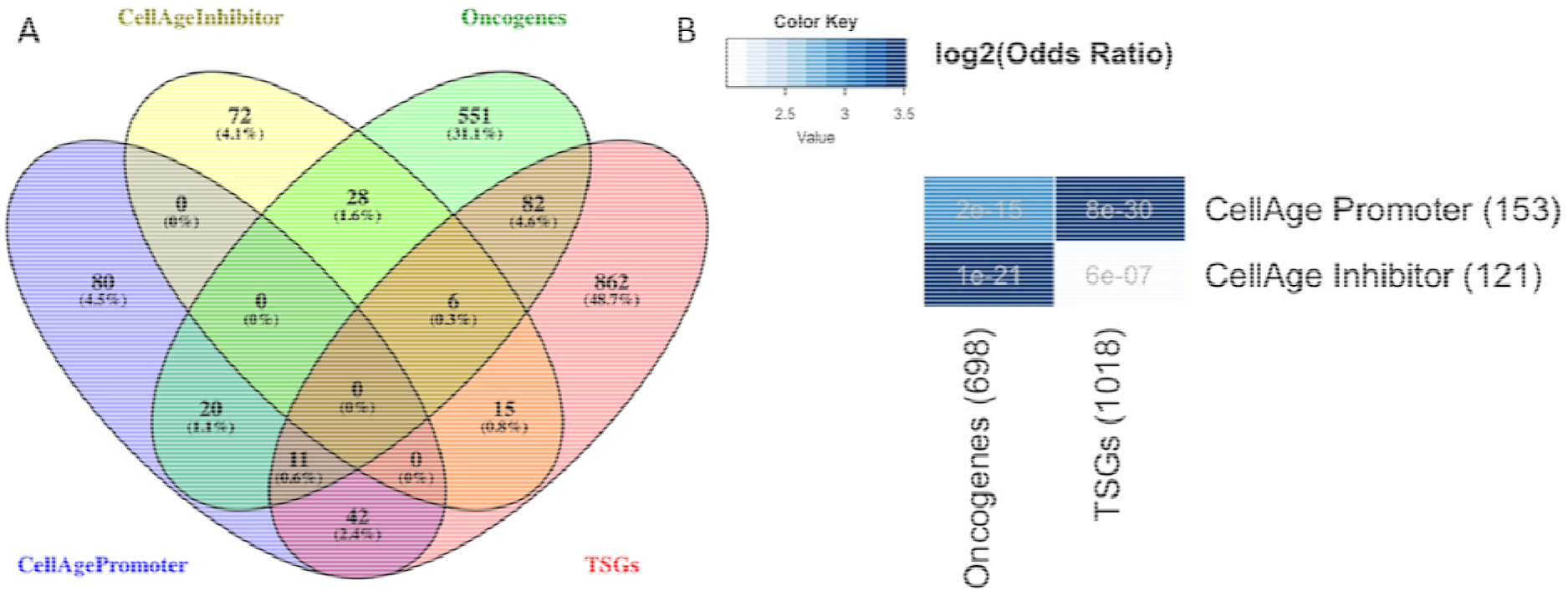
**(A) Overlap between CellAge promoters and inhibitors, and oncogenes and tumour-suppressing genes. (B) Adjusted p-value and odds ratio of the overlap analysis.** The number of overlapping genes in each category was significant (p<0.05, Fisher’s exact test with BH correction). P-values are shown in grey writing for each comparison. Data available in SI Table 23 – 26.

The significant overlap between CellAge genes and cancer indicates a close relationship between both processes. Specifically, the overlap between CellAge inhibitors and oncogenes, and the overlap between CellAge promoters and TSGs were more significant, with lower p-values and larger odds ratios (Figure 4) (Oliveros, 2015).

Gene ontology (GO) enrichment analyses was performed using WebGestalt to identify the function of the overlapping genes (Wang et al., 2017). Overlapping genes between CellAge senescence promoters and TSGs were enriched in GO terms related to p53 signalling and cell cycle phase transition (SI Figure 7A). The enriched functions of overlapping genes between CellAge senescence promoters and oncogenes were mainly related to immune system processes and response to stress (SI Figure 7B). Overlapping genes between CellAge senescence inhibitors and TSGs were enriched in only 5 terms, which are cellular response to oxygen-containing compound, positive regulation of chromatin organization, and terms relating to female sex differentiation (SI Figure 7C). Finally, overlapping genes between CellAge senescence inhibitors and oncogenes were related in processes such as negative regulation of nucleic acid-templated transcription, cellular response to stress, and cell proliferation (SI Figure 7D). All of the functional enrichment data can be found in SI Table 27 – 30.

## NETWORK ANALYSIS

The CellAge genes form both protein-protein and gene co-expression networks. The formation of a PPI network is significant in itself given that only ∼4% of the genes in a randomly chosen gene dataset of similar size are interconnected (Tacutu et al., 2011). In order to have a more holistic view of CS, we were interested in the topological parameters of the networks that CS genes form. For this, several types of networks were constructed using the CellAge genes as seeds: the CS protein-protein interaction (PPI) network, along with two CS gene co-expression networks built using RNA-seq and microarray data. Biological networks generally have a scale-free topology in which the majority of genes (nodes) have few interactions (edges), while some have many more interactions, resulting in a power law distribution of the node degree (the number of interactions per node) (Vidal et al., 2011; Wolfson et al., 2009). As expected, the node-degree distribution of the above networks does confirm the scale-free structure (SI Figure 8). SI Table 31 presents the network summary statistics for the resulting networks.

The network parameters we looked at were Degree, Betweenness Centrality (BC), Closeness Centrality (CC), and Increased Connectivity (IC). The degree is the number of interactions per node and nodes with high degree scores are termed network hubs. BC is a measure of the proportion of shortest paths between all node pairs in the network that cross the node in question. The nodes with high BC are network bottlenecks and may connect large portions of the network which would not otherwise communicate effectively or may monitor information flow from disparate regions in the network (Vidal et al., 2011). CC is a measure of how close a certain node is to all other nodes and is calculated with the inverse of the sum of shortest paths to all other nodes. Lower CC scores indicate that nodes are more central to the network, while high CC scores indicate the node may be on the periphery of the network and thus less central. The IC for each node measures the statistical significance for any over-representation of interactions between a given node and a specific subset of nodes (in our case CellAge proteins) when compared to what is expected by chance. Taken together, genes that score highly for degree, BC, CC, and IC within the senescence networks are likely important regulators of CS even if up until now they have not been identified as CS genes.

Looking at the topology of CS networks, the PPI network, microarray-based co-expression network, and RNA-seq co-expression network possess comparable scale-free structures. However, gene co-expression data is less influenced by publication bias. This is particularly important considering published literature often reports positive protein-protein interactions over protein interactions that do not exist (Gillis et al., 2014). The lack of negative results for protein interaction publications complicates the interpretation of PPI networks even more, as the absence of edges in networks does not necessarily mean they do not exist. On the other hand, RNA-seq and microarray co-expression data, while not influenced by publication bias, does not give indications of actual experimentally demonstrated interactions (physical or genetic). Furthermore, RNA read counts do not directly correlate to protein numbers, with previous studies reporting that only 40% of the variation in protein concentration can be attributed to mRNA levels, an important aspect to keep in mind when interpreting RNA-seq data (Vogel and Marcotte, 2012). Finally, the microarray network was constructed using the COXPRESdb (V6), which contains 73,083 human samples and offered another degree of validation (Okamura et al., 2015). Although RNA-seq reportedly detects more DEGs including ncRNAs (Rao et al., 2018), GeneFriends (van Dam et al., 2015) contains 4,133 human samples, far less than the microarray database from COXPRESdb.

### The Protein-Protein Interaction Network Associated with CS

We only used interactions from human proteins to build the CellAge PPI network. The network was built by taking the CS-associated genes, their first order partners and the interactions between them from the BioGrid database. Interestingly, a very large portion of CS genes and their partners formed a single large network with 1,661 nodes (further analysed here) and several smaller islands (in total the group of networks amounted for 2,643 nodes).

The genes with the highest degree scores are *TP53*, *HDAC1*, *BRCA1*, *EP300* and *MDM2*. Expectedly, several of these genes also possessed the highest BC: *TP53*, *HDAC1*, *BRCA1* and *MDM2* (with *BAG3*, a gene with a slightly smaller degree also within the top 5). On the other hand, focusing more on specificity, the 5 genes with significant IC with existing CellAge genes were: *CCND2*, *SMAD3*, *EGR1*, *CCND1* and *CDKN2A*. Of note among these nodes, *EP300*, *MDM2*, *CCND2*, *SMAD3* and *EGR1* were not already present in CellAge. SI Figure 9 summarises the gene intersection across the computed network parameters, whilst SI Table 32 identifies potential senescence regulators not already present in CellAge from the PPI network. We found that from the top 12 PPI candidates, 11 have been recently shown to regulate senescence in human cell lines and will be added to CellAge build 2 (SI Table 32).

Using DAVID, we found that the following terms were enriched across the 1,661 genes in the PPI network: Transcription, DNA damage & repair, Proteasome & ubiquitin, cell cycle, and ATP pathway (Huang da et al., 2009a, b) (SI Table 33). These results are all in line with previously described hallmarks of cellular senescence (Hernandez-Segura et al., 2018).

It is prudent to note that centrality measures in PPI networks must be interpreted with caution due to publication bias that can be an inherent part of the network (Safari-Alighiarloo et al., 2016; Sanz-Pamplona et al., 2012). The top network genes identified from the PPI network are likely to be heavily influenced by publication bias (Reguly et al., 2006). Looking at the average PubMed hits of the gene symbol in the title or abstract revealed a mean result count of approximately 3,073 per gene, far higher than the genome average (136) or existing CellAge genes (712) (SI Figure 10).

### Unweighted RNA-Seq Co-expression Network

We used CellAge genes that promote and inhibit CS and their co-expressing partners to build a cellular senescence co-expression network. The network consists of a main connected network with 3,198 nodes, and a number of smaller ‘islands’ that are not connected to the main network (Figure 5).

**Figure 5.**
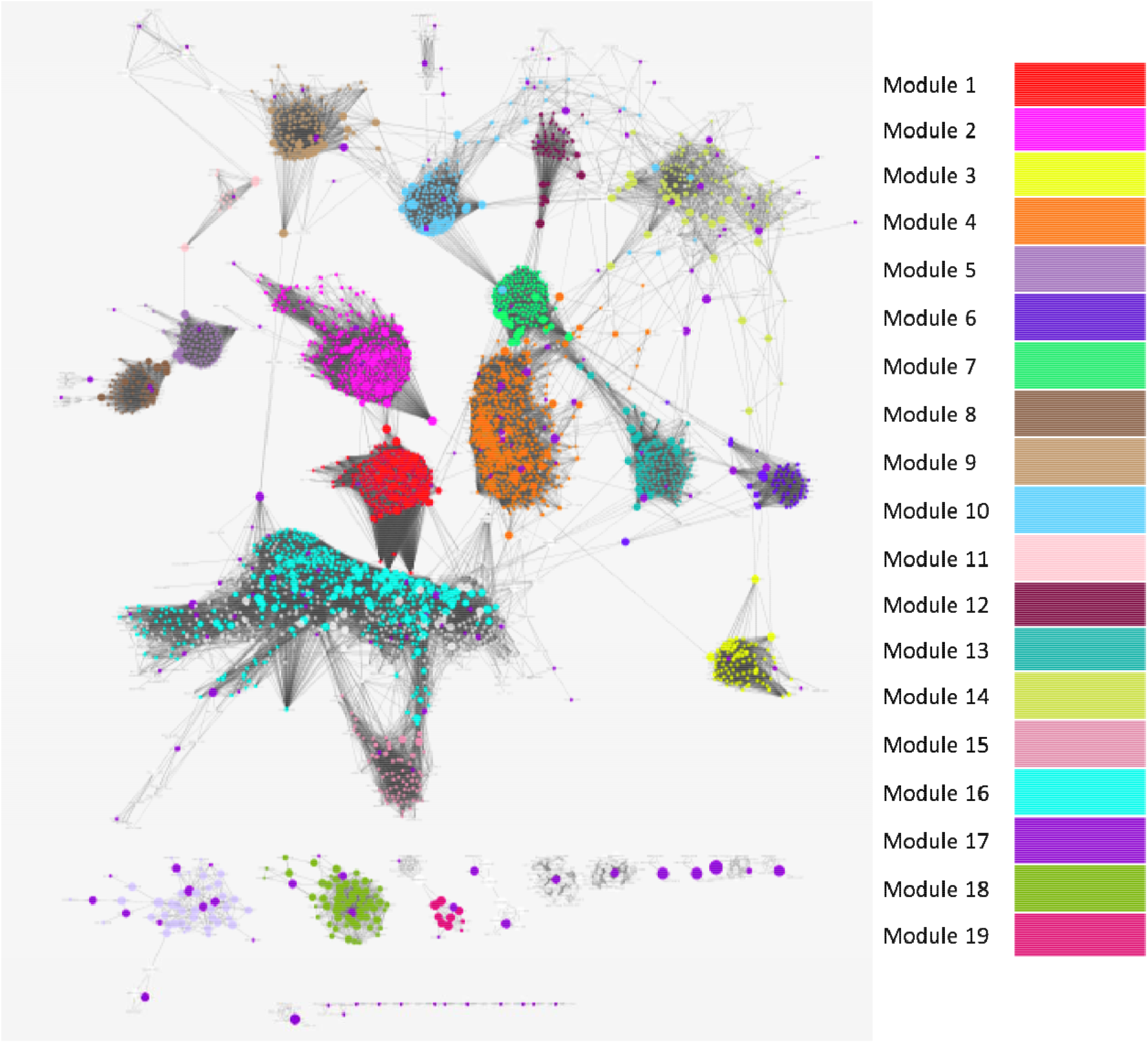
Cluster analysis of the RNA-Seq Unweighted Co-expression Network. The 171 seed nodes obtained from CellAge and their first order interactors. The colours represent the breakdown of the network into clusters. The algorithm revealed 52 distinct clusters, of which we colour and order the 19 clusters with the best rankings for modularity, or in the case of module 17-19, size. The CellAge nodes are coloured in dark purple, appearing throughout the network. Larger nodes have higher betweenness centrality. In order of decreasing modularity, the main function clusters of the modules were related to; Spermatogenesis (Module 1), Synapse (Module 2), Cardiac muscle contraction (Module 3), Cell Cycle (Module 4), Secreted (Module 5), Tudor domain (Module 6), ATP-binding (Module 7), Symport (Sodium ion transport) (Module 8), DNA damage and repair (Module 9), transit peptide:Mitochondrion (Module 10), Steroid metabolism (Module 11), Transcription regulation (Module 12), Protein transport (Module 13), Mitochondrion (Module 14), Heme biosynthesis (Module 15), Innate immunity (Module 16), Signal peptide (Module 17), Keratinocyte (Module 18), Transcription repression (Module 19) (Enrichment results in SI Table 34, genes in SI Table 35).

The main inter-connected network included 130 CellAge genes. Among these, we also found that 14% of them are also human ageing-related genes, reported in GenAge -Human dataset, whereas the remainder of the smaller networks only comprised of 1.6% longevity genes (de Magalhaes et al., 2009). Next, we looked at a number of centrality parameters to see how CellAge genes are characterized compared to the entire network. CellAge genes had a mean BC of 0.00363, whereas the remainder of the genes had a BC of 0.00178, revealing that if CellAge genes are removed, modules within the network may become disconnected more easily. While nodes scoring highly for BC in PPI networks are likely bottleneck regulators of gene expression, this is not necessarily true for co-expression networks. In this case, nodes can also have high BC scores if they are co-activated via various signalling pathways. Although BC alone is not enough to determine which genes are regulating CS, taking BC into account with other network topological parameters can be a good indicator of gene function. Aside from high BC, CellAge genes also had a lower local clustering coefficient of 0.58, compared to a mean of 0.76 across non-CellAge genes, indicating that locally, CellAge genes connect to other genes less than the average for the network. This can also be seen at the degree level, where CellAge genes averaged only 53 connections, compared to an average of 103 connections in non-CellAge genes. Finally, the mean CC score was not significantly different between CellAge nodes and other genes in the network (0.148 in CellAge vs 0.158). CellAge genes were therefore more likely to be bottlenecks in signalling across different modules and occupy localised areas with lower network redundancy, suggesting that perturbations in their expression might have a greater impact on linking different underlying cellular processes.

The topological analysis of the main network component as a whole revealed a more modular topology than the PPI network, resulting in genes tending not to appear in multiple measures of centrality. There were 23 nodes with significant IC with senescence-related genes, including *PTPN6*, *LAPTM5*, *CORO1A*, *CCNB2* and *HPF1*. No node from the top 5 IC was present in the top 5 genes with high BC, CC, or Degree. Overall, the primary candidates of interest included *KDM4C*, which had a significant IC and was at the top 1% of CC and top 5% of BC, along with *PTPN6*, *SASH3* and *ARHGAP30*, which all had significant IC values and were at the top 5% of BC. We found that *KDM4C* and *PTPN6* have been shown to regulate CS in human cell lines, and will be added to build 2 of CellAge (Sun et al., 2015; Yu et al., 2018).

Previous studies have advocated that measures of centrality are generally important to identify key network components, with BC being one of the most common measures. However, it has also been postulated mathematically that intra-modular BC is more important than inter-modular BC (Langfelder et al., 2013). Therefore, by isolating network clusters of interest and identifying genes with high BC or centrality within submodules, we propose to identify new senescence regulators from the co-expression network.

Using the CytoCluster app (see Networks in Materials and Methods section) (Li et al., 2017), we found 54 clusters in the network, of which we represent the top clusters coloured according to modularity (Module 1-16) or size (Module 17-19) (Figure 5). Reactome pathway enrichment for all main clusters highlighted cell cycle and immune system terms in the two largest clusters (Huang da et al., 2009a, b). The largest cluster of 460 nodes (17 CellAge nodes, Module 4), possessed a high modularity score and was strongly associated with cell cycle genes, including the following general terms: Cell Cycle; Cell Cycle, Mitotic; Mitotic Prometaphase; Resolution of Sister Chromatid Cohesion; and DNA Repair. The second largest cluster (Module 16), however, had weak modularity (ranking 26); it comprised of 450 nodes (19 CellAge nodes) and was enriched for immune-related pathways including: Adaptive Immune System; Innate Immune System; Immunoregulatory interactions between a Lymphoid and a non-Lymphoid cell; Neutrophil degranulation; and Cytokine Signaling in Immune system. Cluster 4 and Cluster 5 were not enriched for Reactome Pathways. A visual inspection showed a number of bottleneck genes between Module 1 and Module 16, consistent with the role of the immune system in clearance and surveillance of senescence cells and the secretion of immunomodulators by senescent cells (Hoenicke and Zender, 2012) (SI Table 34).

We were also interested in visualising areas in the network with a high local clustering coefficient, as this parameter represents areas with many neighbourhood interactions and, therefore, more robust areas in the network. It was found that the two clusters of interest, enriched for cell cycle terms and immune system terms, overlapped with regions of lower clustering coefficient, potentially implying parts of the biological system with less redundancy in the underlying process. Figure 6 depicts regions of high local clustering coefficient in the network (orange) and regions less well connected locally (green).

**Figure 6.**
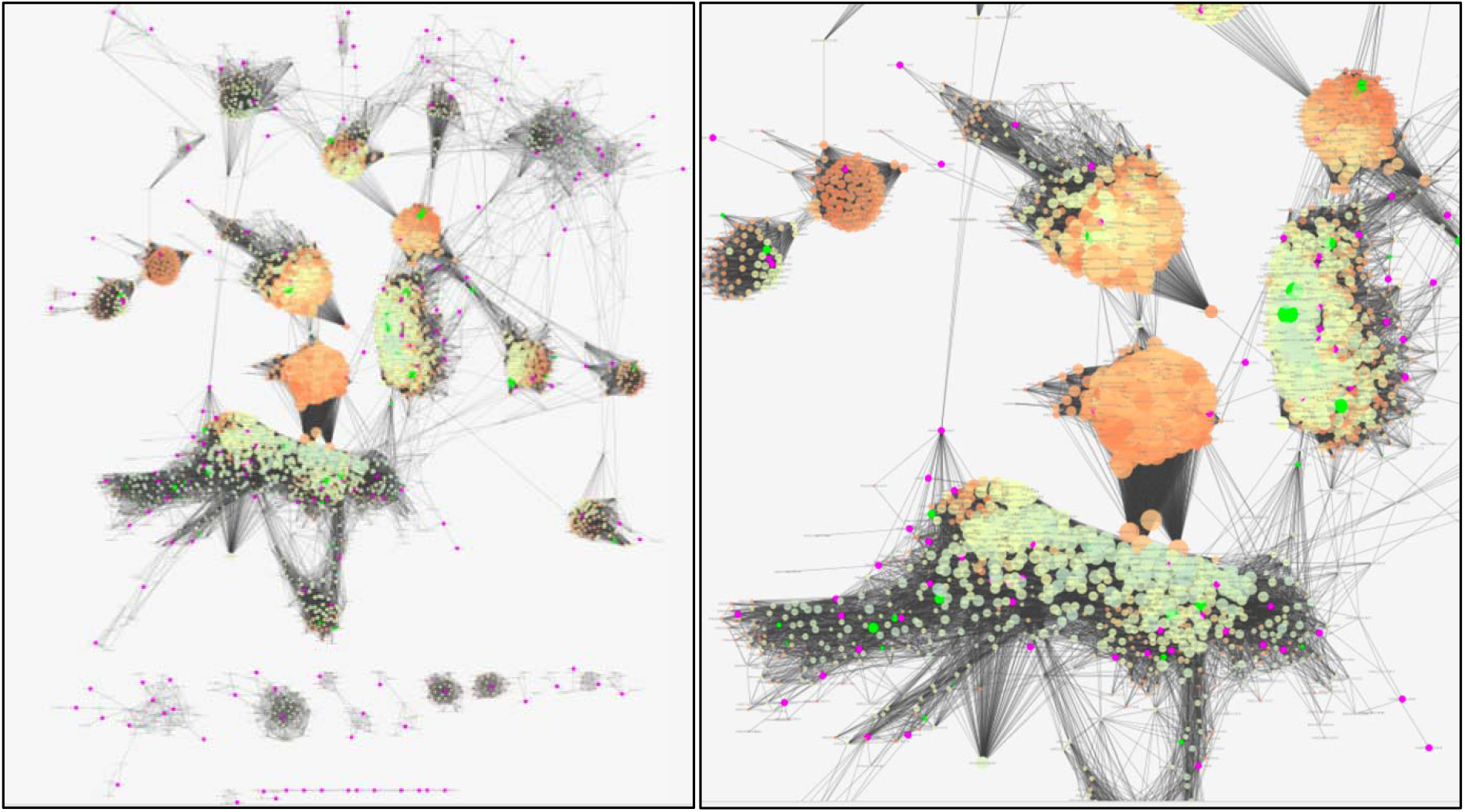
RNA-Seq Unweighted Co-expression Network, local clustering. Red/Orange represents nodes with high clustering coefficient, whereas pale green represents nodes with lower clustering coefficient. Degree is also weighted using node size. CellAge nodes are coloured purple, and GenAge Human nodes are also shown and highlighted in bright green.

### Unweighted Microarray Co-expression Network

We also made an unweighted microarray co-expression network built from the COXPRESdb database of microarray gene co-expression (V6) (Okamura et al., 2015) (SI Figure 11). Compared with the RNA-seq co-expression network, the microarray network is significantly smaller, and only included 34% of the CellAge genes (SI Table 31). However, we found that *SMC4* was an important bottleneck in the microarray network, being in the top 5% CC and IC (SI Figure 11D; 11E). *SMC4* was not independently associated with senescence despite being part of the condensing II complex, which is related to cell senescence (Yokoyama et al., 2015). Furthermore, *SMC4* is associated with cell cycle progression and DNA repair, two key antagonist mechanisms of cell senescence development (d’Adda di Fagagna, 2008; Zhang et al., 2016). *SMC4* has been linked to cell cycle progression, proliferation regulation, and DNA damage repair, in accordance to the most significantly highlighted functional clusters in the module 2 and in the whole Microarray network (SI Figure 12; SI Table 38 and 39) (Dai et al., 2016; Muramatsu et al., 2016). There was limited overlap between the microarray co-expression network and the RNA-seq co-expression network, although this is not surprising considering the higher specificity and sensitivity, and ability to detect low-abundance transcripts in RNA-seq (Wang et al., 2009).

### Experimental Validation of Senescence Candidates

We set out to test if candidate genes from our network analyses are indeed senescence inhibitors using an siRNA-based approach, whereby knockdowns enable the p16 and/or the p21 senescence pathway to be induced, leading to senescence (Stein et al., 1999). In total, we tested a total of 26 potential senescence inhibitor candidates. Of these 26 candidates, 20 were chosen using the HAGR tool GeneFriends, a guilt-by-association database to find co-expressed genes (van Dam et al., 2015). For this, we used the CellAge CS inhibitors as seed genes, with the assumption that genes co-expressed with senescence inhibitors would also inhibit senescence, and generated a list of the top co-expressed genes with CS inhibitors based on RNA-seq data (SI Table 40). Furthermore, CellAge has multiple ways of partitioning genes including the type of senescence the genes are involved in (replicative, oncogenic, and stress-induced). We decided to look for genes co-expressed with stress-induced premature senescence (SIPS) inhibitors as well. We generated a list of genes that are co-expressed with the CellAge SIPS (SI Table 41). A previous paper published a set of gene signatures that are overexpressed and underexpressed in replicative cell senescence (RS) (Chatsirisupachai et al., 2019). We chose to validate five additional genes that were both co-expressed with the CellAge SIPS and are present as underexpressed in our gene signatures in RS. Finally, we chose *SMC4* from the microarray network due to its interaction with other senescence genes within the network, its association with cell cycle progression, and the fact that it is underexpressed in senescent cells, indicating it may be inhibiting senescence in replicating cells. The genes chosen, along with experimental validation results are shown in Figure 7, while the justification for our validation and Z-scores are shown in SI Table 42 and 43 respectively.

**Figure 7.**
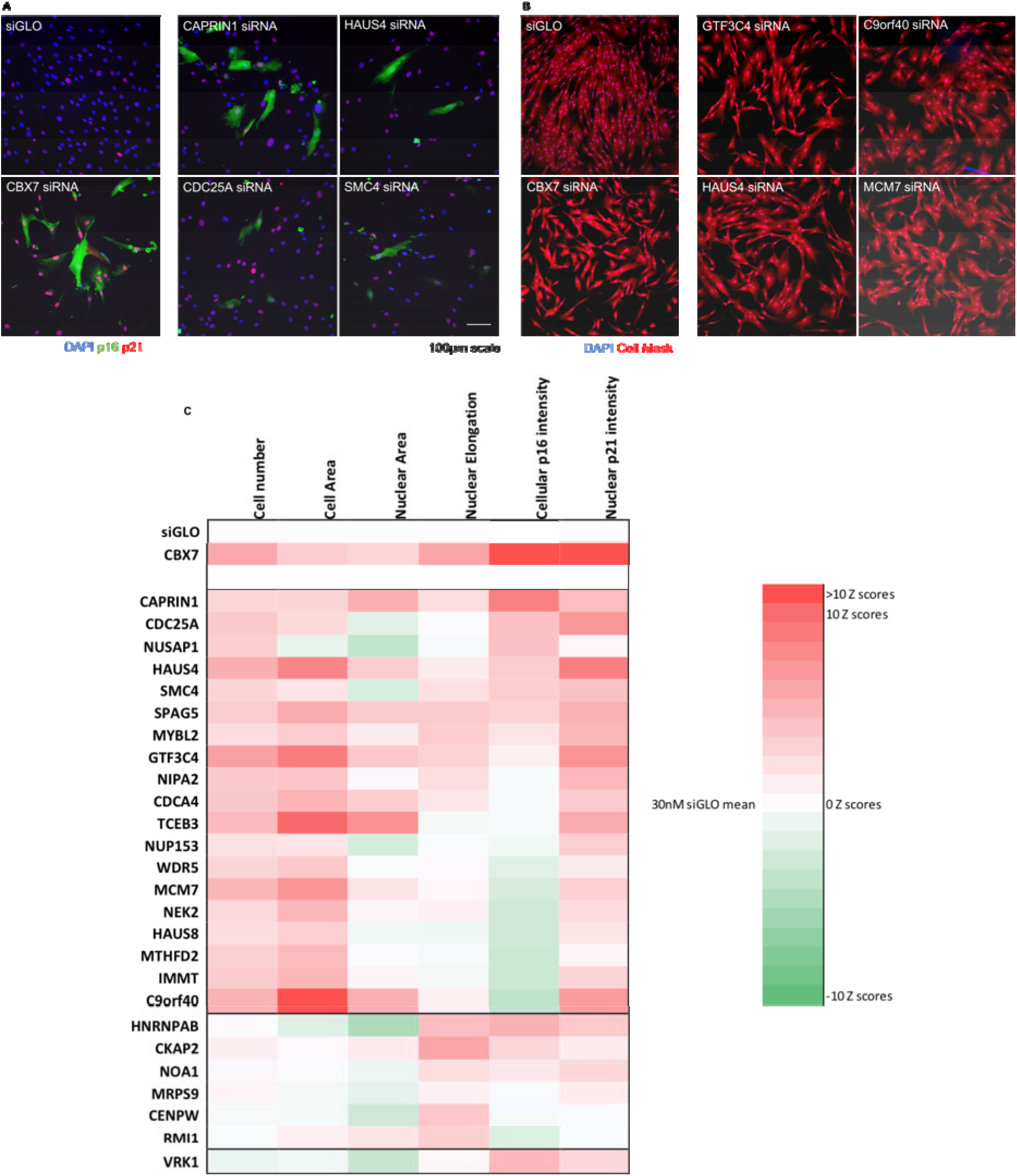
Experimental validation of 26 senescence candidates. **(A) Fibroblasts stained with DAPI (blue), p16 (green) and p21 (red) following transfection with negative control siRNA (siGLO), positive control siRNA (CBX7), or CAPRIN1, HAUS4, CDC25A and SMC4 siRNA. Size bar, 100µm. **(B)** Fibroblasts stained with DAPI (blue) and Cell Mask (red) following transfection with negative control siRNA (siGLO), positive control siRNA (CBX7), or GTF3C4, C9orf40, HAUS4, and MCM7 siRNA.** Size bar, 100µm. **(C) Multi-parameter analysis of cellular and nuclear senescence-associated morphological measures, cellular p16 intensity and nuclear p21 intensity.** Colour coding using to illustrate the number of Z-scores of the experimental siRNA from the siGLO negative control mean. Red values indicate Z-scores closer to the CBX7 positive senescence control. One independent experiment with six technical replicates. All Z-scores available in SI Table 43.

Senescent cells, including fibroblasts, can be characterised by a panel of senescence markers. Markers include a decrease in cell number along with morphological measures such as an increase in cell and nuclear area, and nuclear elongation. Furthermore, the senescence phenotype involves an increase in p16 expression, which is a tumour suppressor protein, along with an increase in p21 expression, a regulator of the cell cycle (Stein et al., 1999). We silenced *CBX7*, a potent senescence inhibitor, as a positive control for senescence induction (Gil et al., 2004). We performed a transient siRNA transfection for 26 candidates and identified those siRNAs which generated the induction of a senescence phenotype compared to the siGLO negative control using multiparameter analysis (Kuilman et al., 2010). Of the 26 genes tested, over 70% (n=19) induced a decrease in cell number greater than 1 Z score, and of these, only one gene did not induce at least one phenotypic change towards senescent morphology (i.e. direction of change similar to the *CBX7* siRNA positive control, Figure 7; SI Table 43). 23% of the candidates (n=6) were top hits, activating both the p16 and p21 pathways, decreasing cell number, and altering at least one phenotype towards a senescent morphology. 46% of the siRNAs (n=12) altered at least one phenotype towards senescent morphology and increased mean p21 intensity but did not increase mean p16 intensity. Finally, one of the siRNAs increased p16 intensity, although there was no increase in p21 intensity or any phenotypic changes towards senescence morphology.

Six of the 26 genes tested had no significant effect on cell number, although five of the genes had one or more phenotypic changes towards senescent morphology. *VRK1* siRNA was the only case which increased cell number and decreased nuclear area greater than 1 Z score (i.e. towards a more proliferative morphology). In general, we have shown the power of networks in predicting gene function, with 18 genes inducing phenotypes towards senescence (69%) and six top hits which decreased cell number, altered at least one phenotype towards senescent morphology, increased mean cellular p16 intensity (p16), and increased mean nuclear p21 intensity (p21) (*CAPRIN1*, *CDC25A*, *HAUS4*, *SMC4*, *SPAG5*, *MYBL2*) (23%).

## DISCUSSION

CellAge aims to be the benchmark database of genes associated with cellular senescence and we expect it to be an important new resource for the scientific community. The development of CellAge has also provided us with the means to perform systematic analyses of CS. While showcasing the functionality of CellAge in this manuscript, we have also explored the links between CS and ageing, ARDs, and cancer. At the same time, we have aimed to expand the knowledge on both the evolution and function of senescence genes, and on how CS genes interact and form genetic networks. We showed that the use of CellAge may help in identifying new senescence-related genes and we have validated several such genes experimentally. As the body of knowledge around senescence grows, it is our aim to maintain a quality resource to allow integrative analyses and guide future experiments. We began our CellAge analysis by gaining further insight into the function of CellAge genes. Unsurprisingly, promoters of CS were enriched for both VEGF and TNF signalling (SI Figure 3; SI Table 8 and 9). Secretion of VEGF is a component of the senescence phenotype and has been shown to contribute towards cancer progression (Coppe et al., 2006). Interestingly, the CellAge genes are more strongly conserved in mammals compared to protein-coding genes, an effect not seen in worms, yeast, or flies (SI Figure 1; SI Table 3). Given the role that many of the senescence genes in CellAge play in regulating the cell cycle, it makes sense that they are evolutionarily conserved; it is not entirely surprising that there is a greater evolutionary pressure towards conserving cell cycle tumour suppressor genes than there is towards conserving other genes. Notably, the pattern of evolutionary conservation of CS genes was found to be almost identical to that of cancer-associated genes, apparently reflecting the co-evolution between these two phenomena (Tacutu et al., 2011). Nonetheless, evolutionary genomics in a comparative context allows us to have a more comprehensive understanding of the genetic bases in important phenotypic traits, like longevity (Doherty and de Magalhaes, 2016). During their evolutionary history, it is possible that long-lived species found ways to more efficiently solve problems related to the ageing process (de Magalhaes et al., 2007; Gorbunova et al., 2014). Lineages where naturally important gene regulators (e.g. TP53) have alternative molecular variants or have been lost from their genomes (Belyi et al., 2010; Wichmann et al., 2016) can be investigated as natural knockouts (Albertson et al., 2009), since they have found a different way to solve ageing-related diseases like cancer (Hanahan and Weinberg, 2011; Stearns et al., 2010).

The relationship between CS and longevity was highlighted across various sections of this manuscript. The promoters of senescence were significantly overrepresented in the anti-longevity human orthologues, while the inhibitors of senescence were even more overrepresented in the pro-longevity human orthologues (SI Table 6) (Fernandes et al., 2016). Furthermore, both the CellAge regulators of CS and the overexpressed signatures of CS were significantly overrepresented in the overexpressed ageing signatures from the human, rat, and mouse ageing signature meta-analysis (Palmer et al., manuscript in preparation). Interestingly, we found that the overexpressed signatures of CS overexpressed with age were significantly enriched for leukocyte activation, cell proliferation, and ageing (SI Figure 3B; SI Table 9). The SASP is a known inducer of chronic inflammation in aged tissue (Acosta et al., 2013; van Deursen, 2014), and the enrichment of terms relating to leukocyte activation highlights the role CS plays in activating the immune system via inflammatory factors with age. One tissue that consistently showed different CS expression patterns with age was the uterus. This observations was already noted in a previous study which also observed that DEGs downregulated in cancer were upregulated with age and DEGs upregulated in cancer were downregulated with age in six tissues, but not in the uterus (Chatsirisupachai et al., 2019).

CS genes are not expressed in a tissue-specific manner (SI Figure 4; SI Table 10) and less than half of the CS genes undergo a significant change in expression with age (Figure 2; SI Figure 5), suggesting that the pathways triggering differential expression of CS genes with age are shared between cells across tissues and are under similar genetic controls. Indeed, we found that *CDKN2A* was overexpressed in 19 human tissues with age, albeit only significantly so in 10 of the tissues (SI Table (Chatsirisupachai et al., 2019). Nonetheless, across 10,000 simulated gene expression overlaps, CS genes significantly overexpressing across multiple tissues with age by chance never exceeded seven tissues, highlighting the significant methodical upregulation of CellAge genes and overexpressed signatures of CS like *CDKN2A*, *SPATA18*, and *GDF15*, which all exceeded the maximum number of overlaps in the simulations (SI Table 16, SI Table 18). These genes are potentially contributing towards the upregulation of other CS genes in aged human tissue.

It is prudent to note that ∼60% of the CellAge database was compiled using experiments in human fibroblast cell lines. Of the 20 studies used to compile the signatures of CS, 10 also performed gene manipulation experiments on fibroblasts (Chatsirisupachai et al., 2019). Fibroblasts are present in connective tissues found between other tissue types across the human body, and the tissue samples analysed to compile GTEx likely contained fibroblast gene expression. This may partially explain the lack of tissue-specific CellAge genes. It is further unclear whether the trends in differential expression of the CellAge genes we see across aged human tissue samples is a result of fibroblast senescence, or if heterogenous gene populations are undergoing CS. Single cell RNA-seq has previously been used to elucidate DEGs in heterogenous cell populations of diseased and healthy spleen samples (Jaitin et al., 2014), and applying similar approaches to young and old tissue samples could clarify both which cells are undergoing CS in ageing and whether the CS process varies across different cell types.

We found a strong association between senescence and neoplastic diseases (SI Table 20). This is not surprising given the known role of senescence in tumour suppression. Some CS genes were also shared between many of the ARD classes. These results are in line with a previous analysis investigating the relationship between CS and ARD genes carried out using different datasets (Tacutu et al., 2011). Tacutu *et al* reported significant overlaps (i.e. 138 genes – 53% – in common between CS and cancer vs 21 – 8% – between CellAge and neoplasms); many more than we did. The study found that many genes shared between CS and several non-cancer ARDs are also involved in cancer. While removing cancer genes from our ARD dataset did not result in such a striking effect, it nonetheless substantially cut the number of overlaps to a statistically insignificant level, adding weight to the hypothesis that cancer genes have a bridging role between CS and ARDs. Furthermore, we found a significant overlap between both the CellAge inhibitors and promoters of senescence, and oncogenes and TSG (Figure 4). Genes that promote senescence, however, tended to be tumour suppressors, while genes that inhibit senescence tended to be oncogenes, a finding that is consistent with the classical view of cellular senescence as a tumour suppressor mechanism.

We next explored what information could be obtained by applying a network analysis to CellAge. From the list of CellAge genes, three networks of CS were generated: a PPI network and two co-expression networks, with the aim of identifying new senescence regulators based primarily on network centrality of the genes.

The examination of the PPI network to identify possible regulators based on centrality and connectivity to existing CS genes revealed 12 central genes in the network (SI Table 32). A further 10 genes were identified by the same criteria, but these were already recorded in the CellAge database. We looked at the RNA-Seq co-expression network in detail, using the main connected component of 3,198 genes to find highly central genes to the network as a whole, and those occupying subnetworks of interest. The RNA-Seq was a highly modular network, separated into some subnetworks of distinct functions (Figure 5). The two largest and more central networks contained a number of known senescence genes. We expanded the analysis of these networks in particular, identifying a number of bottleneck nodes. Cluster 1 was enriched for cell cycle processes, which is not overly surprising given that senescence involves changes in cell cycle progression. However, cluster 2 comprised of enriched terms relating to immune system function. One of the aims in biogerontology is to understand and reverse the effects of ageing on the immune system. SI Table 37 highlights the genes in both clusters that are potential CS bottlenecks within the network and may warrant further study.

Using siRNAs, we were able to test the potential role of 26 gene candidates in inhibiting senescence (Figure 7). The list of candidates was primarily compiled using CellAge inhibitors as seeds to generate co-expressed genes in GeneFriends, a collection of RNA-seq co-expression data (van Dam et al., 2015) (SI Table 42). Of the 26 genes, 6 were top hits, causing cells to undergo morphological changes towards senescent phenotypes, decreasing overall cell number, and activating both the p16 and p21 pathways. A further 13 siRNAs also induced a decrease in cell number, altered at least one phenotype towards senescent morphology, and increased mean p21 or p16 intensity, but not both. SI Table 44 highlights the four CS candidates we found that have not yet been associated with senescence. We have showcased how co-expression networks can be used to accurately infer senescence gene candidates, which can then be experimentally verified.

Cellular senescence is one of the hallmarks of ageing (Lopez-Otin et al., 2013) and the accumulation of senescent cells in human tissue with age is implicated as a driver of ageing-related diseases. Indeed, pharmacological approaches targeting senescent cells, like senolytics, are a major and timely area of research that could result in human clinical applications (de Magalhaes and Passos, 2018; de Magalhaes et al., 2017). It is imperative that we fully understand and deconstruct cellular senescence in order to target ageing-related diseases. We hope that CellAge will help researchers understand the role that CS plays in ageing and ageing-related diseases and contributes to the development of drugs and strategies to ameliorate the detrimental effects of senescent cells.

## MATERIALS AND METHODS

### CellAge Compilation

CellAge was compiled following a scientific literature search, manual curation, and annotation, with genes being appended to the database if they met the following criteria:

- Only gene manipulation experiments (gene knockout, gene knockdown, partial or full loss-of-function mutations, over-expression or drug-modulation) were used to identify the role of the genes in cellular senescence. The search focussed on genes from genetic manipulation experiments to ensure objectivity in the selection process.
- The genetic manipulation caused cells to promote or inhibit the CS process in the lab. Cellular senescence was detected by growth arrest, increased SA-β-galactosidase activity, SA-heterochromatin foci, a decrease in BrdU incorporation, changes in morphology, and/or specific gene expression signatures.
- The experiments were performed in primary, immortalized, or cancer human cell lines.

Over half of the experiments were conducted in lung and foreskin fibroblasts. The data was compiled from 230 references. The curated database comprises cell senescence genes together with a number of additional annotations useful in understanding the context of each identified CS gene (SI Table 45).

We categorised genes according to three types of senescence: replicative, oncogene-induced or stress-induced. We also recorded whether a gene promotes or inhibits CS. For example, a gene whose overexpression is associated with increased senescence is classified with the ‘promotes’ tag, whereas if the overexpression of a gene inhibits senescence, then it is classified with the ‘inhibits’ tag. Similarly, if the knockout or knockdown of a gene promotes senescence, then it is recorded with the ‘inhibits’ tag. Together with the annotations identified in SI Table 45, we also incorporated a number of secondary annotations into the database such as various gene identifiers, the gene description, gene interaction(s), and quick links to each senescence gene. The CellAge database also provides crosslinks to genes in other HAGR resources i.e. GenAge, GenDR and LongevityMap, which we hope will enable inferences to be made regarding the link between human ageing and CS.

### CellAge Data Sources

Build 1 of CellAge resulted in a total of 279 curated cell senescence genes which we have incorporated into the HAGR suite of ageing resources. The HAGR platform comprises a suite of ageing databases and analysis scripts. The CellAge interface has been designed with the help of JavaScript libraries to enable more efficient retrieval and combinatorial searches of genes. As with the other HAGR databases, we have used PHP to serve the data via an Apache web server. The raw data can be downloaded via the main HAGR downloads page in CSV format or filtered and downloaded from the main search page.

The first part of our work consisted in finding which CS-associated genes are also associated with ARDs or with longevity, using the following data sources:

- Human genes associated with CS: <u>CellAge</u> build 1.
- Human genes associated with human ageing: GenAge human build 19.
- Human orthologues of model organisms’ genes associated with longevity: proOrthologuesPub.tsv and antiOrthologuesPub.tsv file (https://github.com/maglab/genage-analysis/blob/master/Dataset_4_aging_genes.zip) (Fernandes et al., 2016).
- oncogenes: Oncogene database (http://ongene.bioinfo-minzhao.org/index.html)
- tumour suppressor gene database: TSGene 2.0 (https://bioinfo.uth.edu/TSGene/index.html)
- genes associated with ARDs (https://github.com/maglab/genage-analysis/blob/master/Dataset_5_disease_genes.zip) (Fernandes et al., 2016). This data concerns the 21 diseases with the highest number of gene associations, plus asthma, a non-ageing-related respiratory system disease used as a control.

### CellAge Data Analysis

Statistical significance was determined by comparing the p-value of overlapping CellAge gene symbols with the different data sources, computed via a hypergeometric distribution and Fisher’s exact test. We used PubMed to understand the relative research focus across the protein coding genome and incorporate this into the analysis to account for publication bias. We used BioMart to obtain approximately 19,310 protein coding genes, then using an R script we queried NCBI for the publication results based on the gene symbol using the following query (Kinsella et al., 2011; R Core Team, 2018):

(“***GENE_SYMBOL***”[Title/Abstract] AND Homo[ORGN]) NOT Review[PTYP]

The GENE_SYMBOL was replaced in the above query by each of the genes in turn. Certain genes were removed as they matched common words and, therefore, skewed the results: *SET*, *SHE*, *PIP*, *KIT*, *CAMP*, *NODAL*, *GC*, *SDS*, *CA2*, *COPE*, *TH*, *CS*, *TG*, *ACE*, *CAD*, *REST*, *HR*, and *MET*. The result was a dataframe in R comprising variables for the ‘gene’ and the ‘hits’. We used the R package called ‘rentrez’ to query PubMed for the result count (Winter, 2018).

### Evolution of CellAge Genes

The percentage of CellAge genes with orthologues in *Rhesus macaque, Rattus norvegicus, Mus musculus, Saccharomyces cerevisiae, Caenorhabditis elegans*, and *Drosophila melanogaster* were found using Biomart version 88 (Kinsella et al., 2011). We also found the total number of human genes with orthologues in the above species using Biomart. Significance was assessed using a two-tailed z-test with BH correction.

The phylogenetic arrangement included twenty-four species representative of major mammalian groups. The genomes were downloaded in CDS FASTA format from Ensembl (http://www.ensembl.org/) and NCBI (https://www.ncbi.nlm.nih.gov/) (SI Table 5). To remove low quality sequences we used the clustering algorithm of CD-HITest version 4.6 (Fu et al., 2012) with a sequence identity threshold of 90% and an alignment coverage control of 80%. The longest transcript per gene was kept using TransDecoder.LongOrfs and TransDecoder.Predict (https://transdecoder.github.io) with default criteria (Haas and Papanicolaou). In order to identify the orthologs of the 279 CellAge human genes in the other 23 mammalian species, the orthology identification analysis was done using OMA standalone software v. 2.3.1 (Altenhoff et al., 2015). This analysis makes strict pairwise sequence comparisons ‘all-against-all,’ minimizing the error in orthology assignment. The orthologous pairs (homologous genes related by speciation events) are clustered into OrthoGroups (OG) (Altenhoff and Dessimoz, 2009); this was done at the Centre for Genomic Research computing cluster (Linux-based) at the University of Liverpool. The time calibrated tree was obtained from TimeTree (http://www.timetree.org/) and the images were downloaded from PhyloPic (http://phylopic.org/).

### Overlap Analysis

We conducted overlap analysis using R to understand how the CellAge genes and signatures of CS were differentially expressed with GenAge, ARD, and cancer genes. We also examined the overlap between CS genes and differentially expressed signatures of ageing (Palmer et al., manuscript in preparation), and genes differentially expressed in various human tissues with age. Fisher’s exact test was used on the contingency tables and significance was assessed by p-values adjusted via Benjamini-Hochberg (BH) correction. For the comparison of genes differentially expressed in at least one tissue with age between the CS genes and the genome, some genes were differentially expressed in opposite directions across numerous tissues (SI Figure 5A). Genes differentially expressed in both directions were added to the overexpressed and underexpressed DEGs in each CS gene list, and to the total number of genes in the genome to compensate for the duplicate gene count (SI Table 13 and 14). Fisher’s exact test was also used to test for significance of tissue-specific CellAge gene expression. Significance of overlap analysis between CellAge and LAGs was computed using a hypergeometric distribution and FDR was corrected using Bonferroni correction. The GeneOverlap package in R was used to test for overlaps between the CellAge promoters and inhibitors of senescence, and the oncogenes and TSGs (Shen and Sinai, 2013). Results for all overlap analyses were plotted using the ggplot2 library (R Core Team, 2018; Wickham, 2016).

### Simulation of CS Gene Expression in Human Ageing

The RNA-seq gene expression data on GTEx was scrambled in such a way that all protein-coding genes in each tissue were assigned a random paired p and log_2_FC value from the original gene expression data of each respective tissue. The randomly sorted gene expression data was then filtered for significance (p<0.05, moderated t-test with BH correction, absolute log_2_FC>log_2_(1.5)) (Chatsirisupachai et al., 2019; Ritchie et al., 2015), and the CellAge accessions were extracted and overlapped across all the simulated expression data in 26 tissues from GTEx. The probability of a CS gene being overexpressed or underexpressed across multiple tissues by chance was calculated across 10,000 simulations.

### Functional Enrichment

The analysis of CellAge included gene functional enrichment of the database. We used DAVID functional clustering (https://david.ncifcrf.gov/) to identify functional categories associated with CellAge (Huang da et al., 2009a, b).

The Overrepresentation Enrichment Analysis (ORA) of biological processes (Gene Ontology database) was done via the WEB-based Gene SeT AnaLysis Toolkit (WebGestalt) (Wang et al., 2017) for the analysis of CellAge senescence regulators and overexpressed signatures of CS overexpressed in the meta-analysis of ageing signatures, and for the CellAge genes overlapping with tumour suppressor and oncogenes. A p-value cutoff of 0.05 was used, and p-values were adjusted using BH correction. Redundant GO terms were removed and the remaining GO terms were grouped into categories based on their function using Reduce + Visualize Gene Ontology (REVIGO) (Supek et al., 2011). Results were then visualised using and the R package treemap (Tennekes, 2017) (SI Figure 7A – D). Venn diagrams to represent gene overlaps were created using Venny (Oliveros, 2015).

### Networks

We used Cytoscape version 3.6.1 to generate networks and R version 3.3.1 to perform aspects of the statistical analysis (R Core Team, 2018; Shannon et al., 2003). The networks were built starting from a list of seed nodes – all genes included in build 1 of CellAge, part of the Human Ageing Genomic Resources (Tacutu et al., 2018). Network propagation was measured using the Cytoscape plugin Diffusion (Carlin et al., 2017).

The analysis of the fit to the scale-free structure was calculated by the Network Analyzer tool of Cytoscape 3.2.1 (Shannon et al., 2003). Network analyzer is a Cytoscape plugin which performs topological analysis on the network and reports the pillar nodes on the network structure based on a series of mathematical parameters (Degree, BC and CC) (SI Figure 8). Network analyzer also calculates the fit of the distribution of the number of edges per node to the power-law distribution. A significant fit to the power law indicates the presence of a scale free structure in the network (Albert et al., 2000; Safari-Alighiarloo et al., 2016). The analysis was applied to the PPI network, the RNAseq Unweighted Co-expression network, and the Microarray Unweighted Co-expression network of cellular senescence. The Network Analyzer tool was also used to calculate BC, CC, and IC in the networks.

### Protein-Protein Interaction Network

The protein-protein interaction network was built from the BioGrid database of physical multi-validated protein interactions (Biology General Repository for Interaction Datasets) version 3.4.160, using CellAge proteins as seed nodes and extracting the proteins encoded by CellAge genes as well as the first order interactors of CellAge proteins (Chatr-Aryamontri et al., 2017). After removing duplicated edges and self-loops, the network consisted of 2,643 nodes and 16,930 edges. The network was constructed and visualised in Cytoscape version 3.6.1. The “CytoCluster” App in Cytoscape was used to identify modules in the network with the following parameters: HC-PIN algorithm; Weak, Threshold = 2.0; ComplexSize Threshold = 1% (Li et al., 2017).

### Unweighted RNA-Seq Co-Expression Network

The RNA-seq co-expression network was built using CellAge data and RNA-Seq co-expression data taken from Genefriends (http://genefriends.org/RNAseq) (van Dam et al., 2015).

The unweighted co-expression network was built applying the method of correlation threshold selection described by Aoki to the GeneFriends database of RNA-Seq co-expression version 3.1 (Aoki et al., 2007). Aoki initially designed this methodology for plant co-expression network analysis, but it has been successfully applied to build human networks (Bartel et al., 2015). The Pearson Correlation Coefficient (PCC) threshold which generated the database of edges with the lowest network density was selected. The network density is the proportion of existing edges out of all possible edges between all nodes. The lower the network density is the more nodes and fewer edges are included in the network. The lower the number of edges, the higher the minimum correlation in expression between each pair of genes represented by the edges. The higher the number of nodes, the higher the portion of nodes from CellAge included, and, therefore, the more representative the network is of the CellAge database. The PCC threshold of 0.65 generated the database of interactions of RNA-Seq co-expression with the lowest network density, 0.01482 (SI Figure 13A). The unweighted RNA-Seq network was generated and visualised in Cytoscape 3.6.1.

### Microarray Co-Expression Network

The microarray co-expression network was generated using the CellAge genes as seed nodes and their direct interactions and edges, derived using the COXPRESdb database of Microarray co-expression (version Hsa-m2.c2-0) (Okamura et al., 2015). PCC threshold of 0.53 created the Microarray database with the lowest network density, 1.006*10^-2^ (SI Figure 13B). The adjustment of the node-degree distribution to the power law distribution had a correlation of 0.900 and an R-squared of 0.456 (SI Figure 8C). The fit to the power law distribution confirmed the scale-free structure of the network.

### Cell Culture and Reagents

Normal human mammary fibroblasts (HMFs) were obtained from reduction mammoplasty tissue of a 16-year old individual, donor 48 (Stampfer et al., 1981). The cells were seeded at 7,500 cells/cm^2^ and maintained in Dulbecco’s Modified Eagles Medium (DMEM) (Life Technologies, UK) supplemented with 10% foetal bovine serum (FBS) (Labtech.com, UK), 2mM L-glutamine (Life Technologies, UK) and 10µg/mL insulin from bovine pancreas (Sigma). All cells were maintained at 37°C/5% CO_2_. All cells were routinely tested for mycoplasma and shown to be negative.

### siRNA Knockdown Experiments

For high-content analysis (HCA), cells were forward transfected with 30nM siRNA pools at a 1:1:1 ratio (Ambion) using Dharmafect 1 (Dharmacon) in 384-well format. Control siRNA targeting cyclophilin B (siGLO, Dharmacon) or Chromobox homolog 7 (CBX7, Ambion) were also included as indicated. Cells were incubated at 37°C/5% CO_2_ and medium changed after 24hr. Cells were then fixed/stained 96hr later and imaged as described below. The siRNA sequences are provided in SI Table 46.

### Z Score Generation

For each of the parameters analysed, significance was defined as one Z score from the negative control mean. Z scores were generated according to the formula below:

Z score = (mean value of one independent experiment for target siRNA with three technical replicates – mean value of one independent experiment for siGLO with six technical replicates)/standard deviation (SD) for siGLO of one independent experiment with six technical replicates.

### Immunofluorescence Microscopy and High Content Analysis

Cells were fixed with 3.7% paraformaldehyde, permeabilised for 15min using 0.1% Triton X and blocked in 0.25% BSA before primary antibody incubations. Primary antibodies used are listed in SI Table 47. Cells were incubated for 2hr at room temperature with the appropriate AlexaFluor-488 or AlexaFluor-546 conjugated antibody (1:500, Invitrogen), DAPI and CellMask Deep Red (Invitrogen). Images were acquired using the IN Cell 2200 automated microscope (GE) and HCA was performed using the IN Cell Developer software (GE).

## SUPPLEMENTARY MATERIAL

Supplementary figures and citations, tables, and FASTA files are available on the Integrative Genomics of Ageing Group CellAge_supplementary GitHub repository (https://github.com/maglab/CellAge_supplementary).

## AUTHOR CONTRIBUTIONS

R.A.A and J.G.O wrote the manuscript. S.S, A.M, and J.P.M selected CS candidates. E.T and C.L.B performed experimental validation of CS genes. R.A.A, J.G.O, R.T, P.B, K.C, E.J, S.S, D.T.M, J.P.M and D.T did bioinformatics analysis. E.T, D.T, R.T, D.B, A.B, V.E.F, C.L.B, and J.P.M edited the manuscript. J.P.M and R.T conceived the project.

## ACKNOWLEDGMENTS

This work was supported by grants from the Wellcome Trust (104978/Z/14/Z and 208375/Z/17/Z), the Leverhulme Trust (RPG-2016-015), LongeCity and the Biotechnology and Biological Sciences Research Council (BB/R014949/1), EU-Romanian Competitiveness Operational Programme (POC-A.1-A.1.1.4-E-2015), the ‘Comisión Nacional de Investigación Científica y Tecnológica (CONICYT) - Chile’ (doctoral studentship N°21170433), and by the Fund in Memory of Dr. Amir Abramovich. ET is funded by the BBSRC (BB/P002579/1). Richard Gregory in the Centre for Genomic Research (CGR) and Ian C. Smith in the Advanced Research Computing at the University of Liverpool helped with the computing resources.

