## Supplementary material for "A Multidimensional Systems Biology Analysis of Cellular Senescence in Ageing and Disease": SI Figures

A

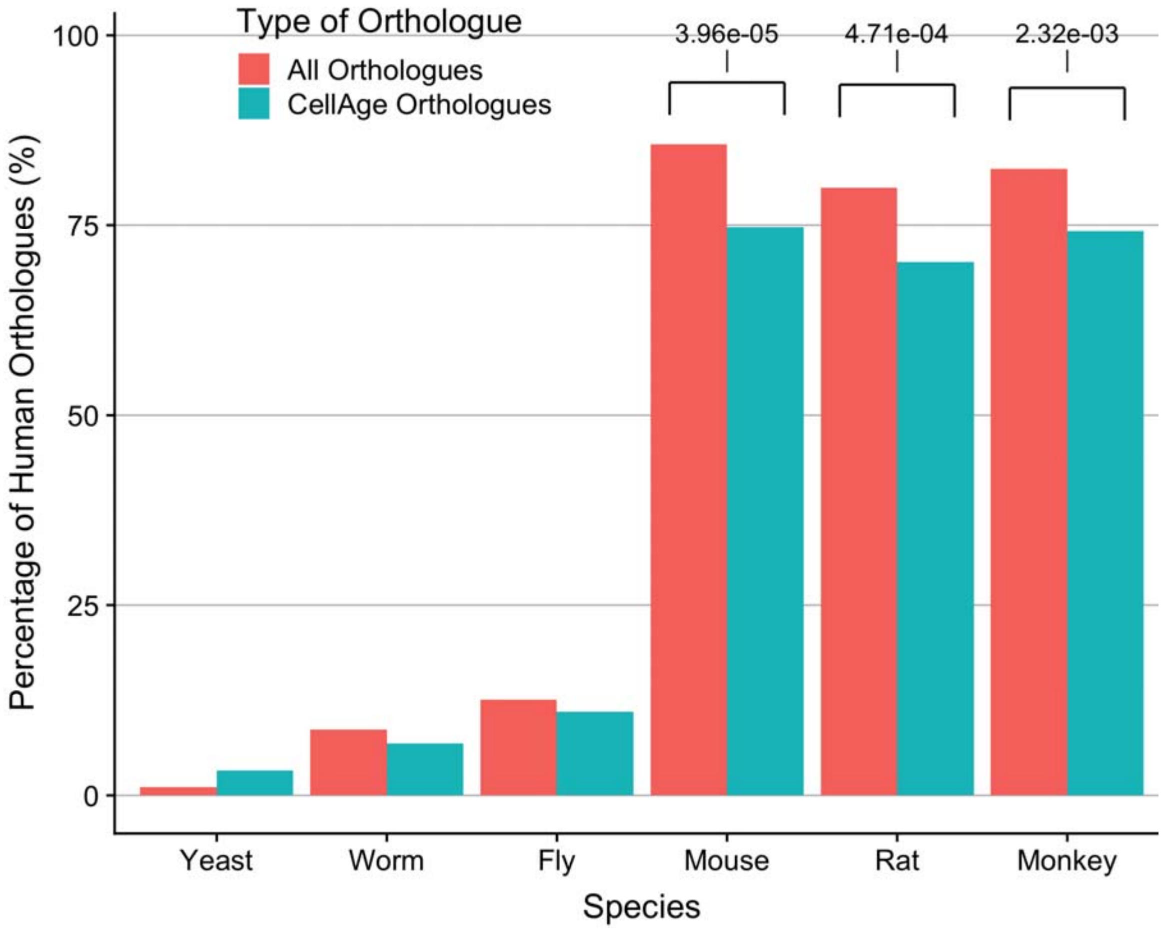
Type of Orthologue
All Orthologues
CellAge Orthologues
3.96e-05
4.71e-04
2.32e-03
Percentage of Human Orthologues (%)
100
75
50
25
0
Yeast
Worm
Fly
Mouse
Rat
Monkey
Species

B

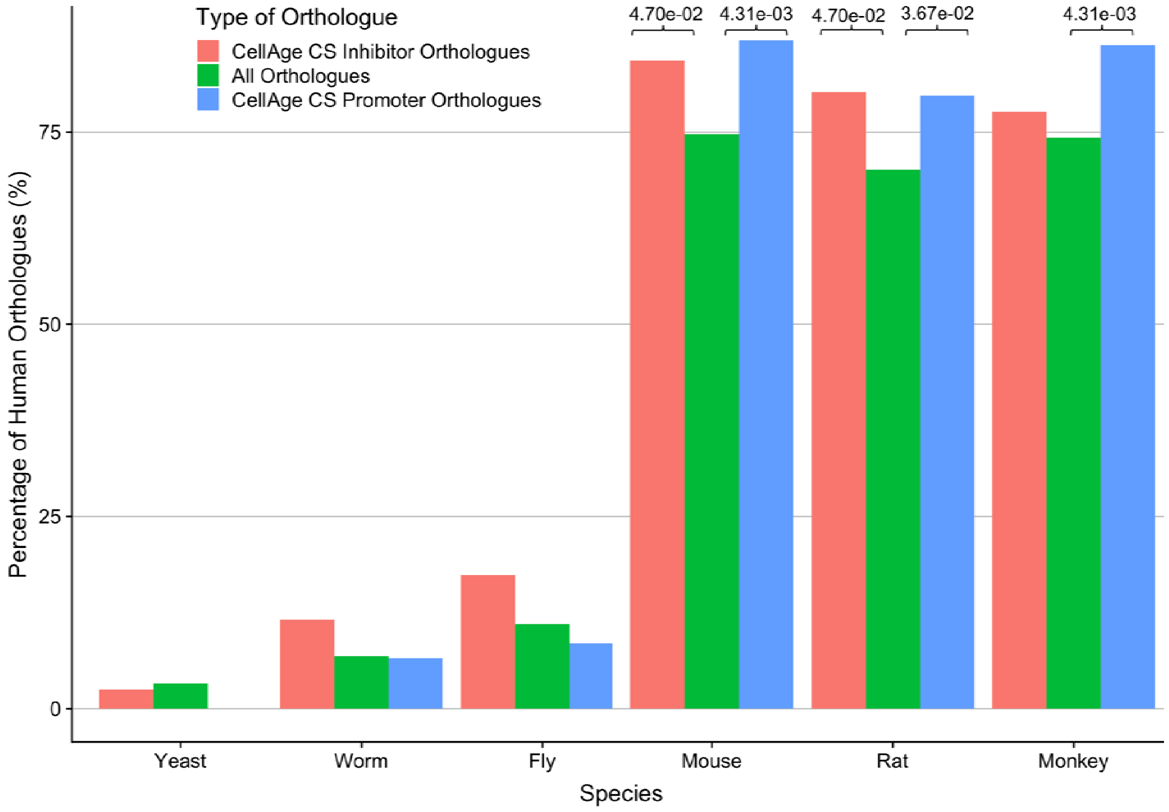
Type of Orthologue
CellAge CS Inhibitor Orthologues
All Orthologues
CellAge CS Promoter Orthologues
4.70e-02
4.31e-03
4.70e-02
3.67e-02
4.31e-03
Percentage of Human Orthologues (%)
75
50
25
0
Yeast
Worm
Fly
Mouse
Rat
Monkey
Species

C

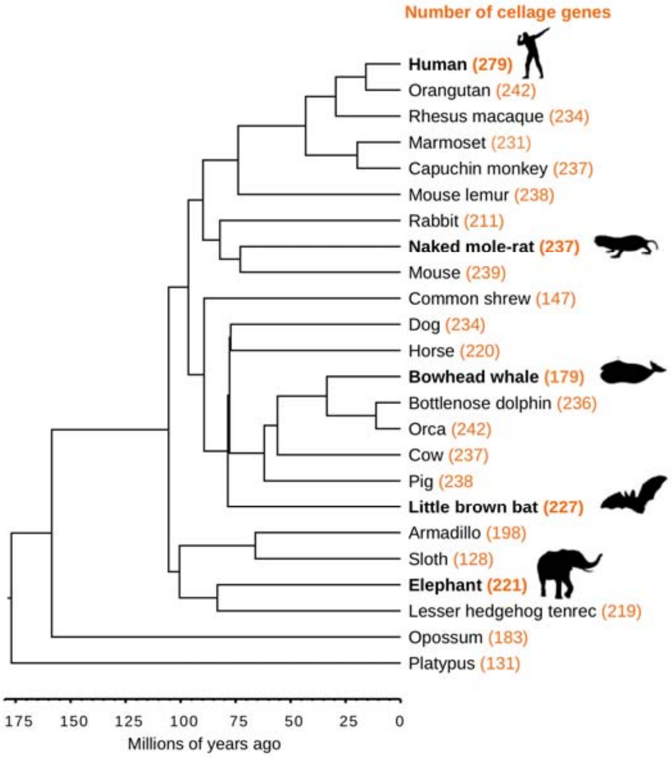


**SI Figure 1. Evolution of the CellAge genes (A) Comparing conservation of the CellAge genes and normal genes in human vs model organisms.** * Indicates significant differences between conservation ($p<0.05$, z-test with BH correction). (**B) Comparing conservation of the CellAge promoters and inhibitors of CS and normal genes in human vs model organisms.** * Indicates significant differences between conservation ($p<0.05$, z-test with BH correction). (**C) Orthologous CellAge genes presents in 24 mammalian species.** The number of orthologues are shown in orange. The species in bold are considered long-lived with medical importance in anti-ageing research.

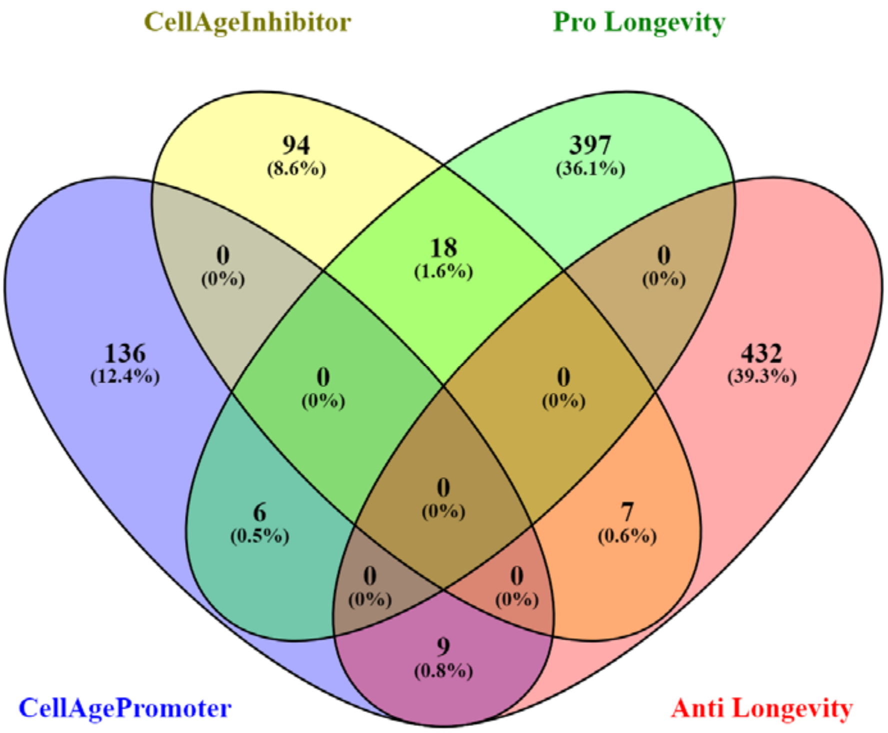


**SI Figure 2. Overlap between CellAge promoters and inhibitors, and pro and anti-longevity genes.**

A

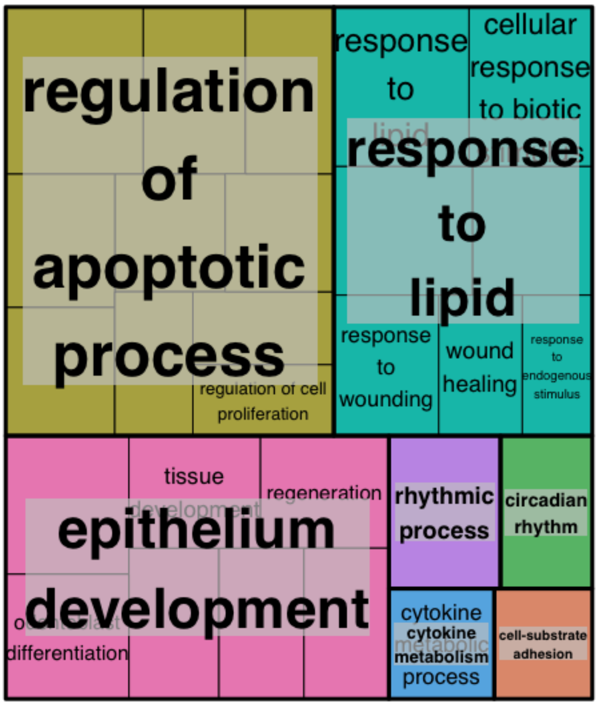

B

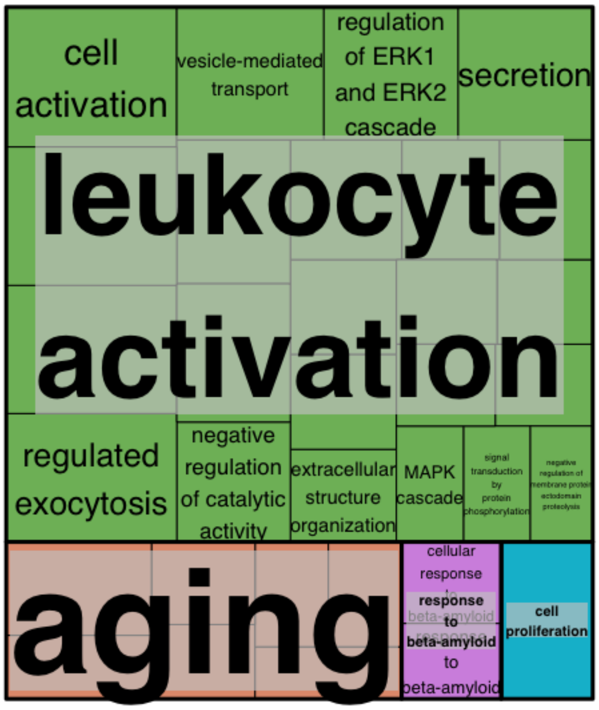


**SI Figure 3. Treemaps of enriched GO terms for (A) CellAge promoters of CS overexpressed with age and (B) overexpressed signatures of CS overexpressed with age.**

A

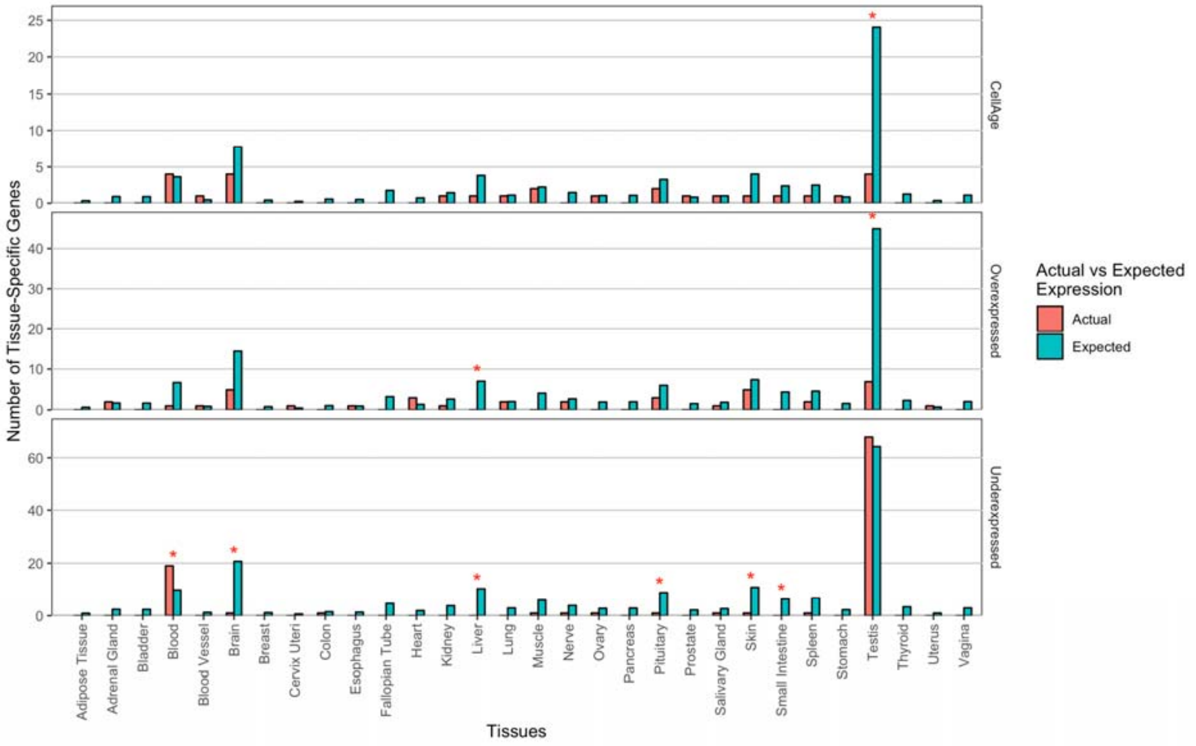

B

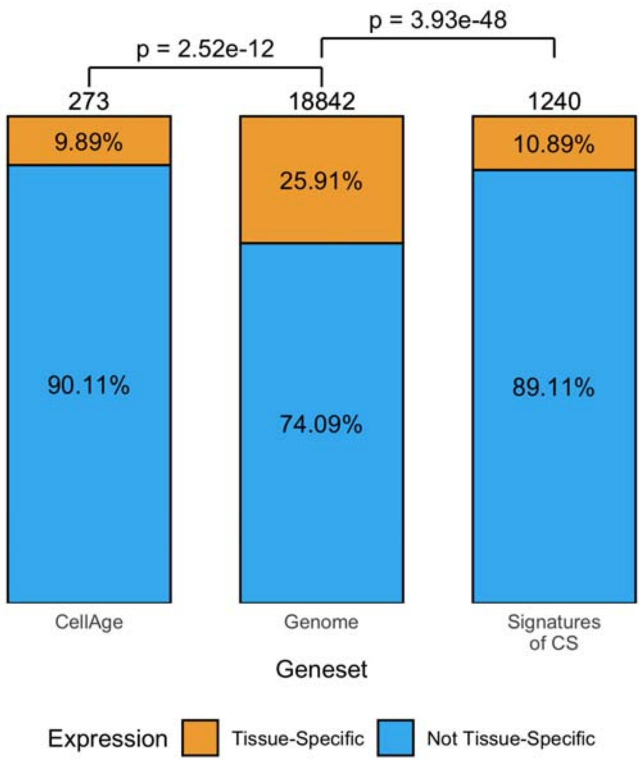


**SI Figure 4. (A) Comparison between the actual number of CS genes expressed in a tissue-specific manner vs. the expected number.** * Represents a significant difference (p<0.05, Fisher's exact test with BH correction). Full data available in SI Table 9. **(B) Comparison between the ratio of tissue-specific to non-tissue-specific genes in the CS datasets vs. all protein-coding genes**. P-values denote significance (p<0.05, Fisher's exact test with BH correction).

A

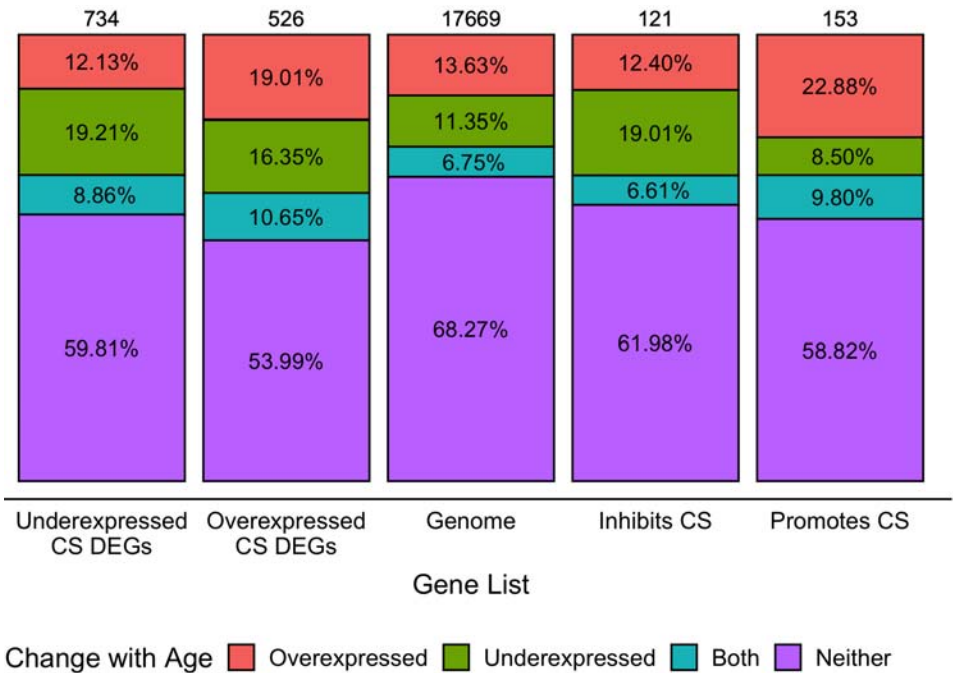

B

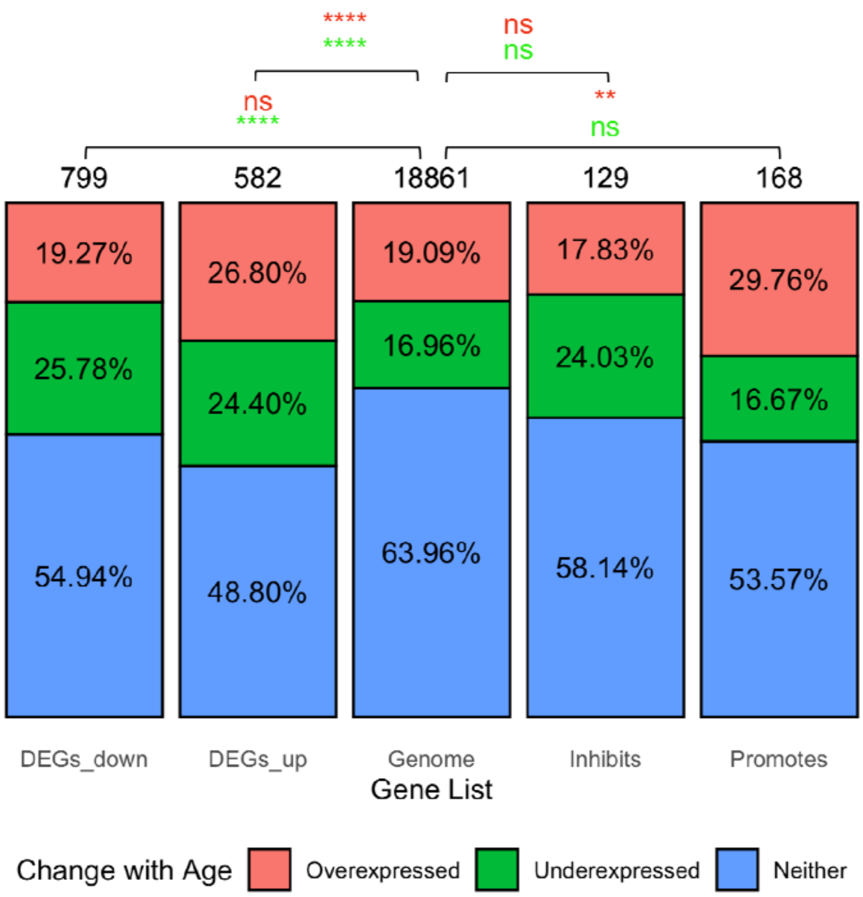


**SI Figure 5. (A) Breakdown of CS genes differentially expressed in at least one tissue with age.** Numbers over each bar represent total number of genes in each dataset. Some genes were differentially expressed in opposite directions with age across multiple tissues, and as such are listed as 'both.' **(B) Comparison between ratio of CS genes differentially expressed in at least one tissue with age vs. all protein-coding genes differentially expressed in at least onet issue with age.** Red values indicate comparison between overexpressed genes, whereas green values indicate comparisons between underexpressed genes. Significance assessed using Fishers exact test with BH correction (NS – not significant, * – $p<0.05$, ** – $p<0.01$, *** – $p<0.001$, **** – $p<0.0001$). Total number of genes differ in A and B because 'both' was added to both overexpressed and underexpressed and to gene totals to compensate.

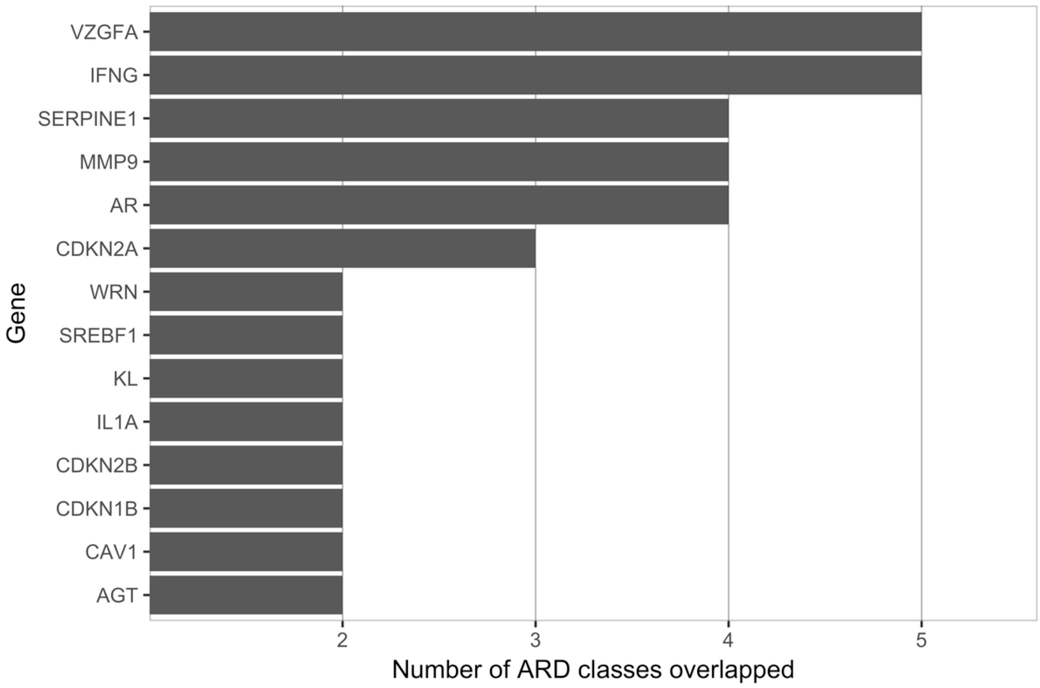


**SI Figure 6. Results of CellAge genes that overlap with multiple ARD classes.**

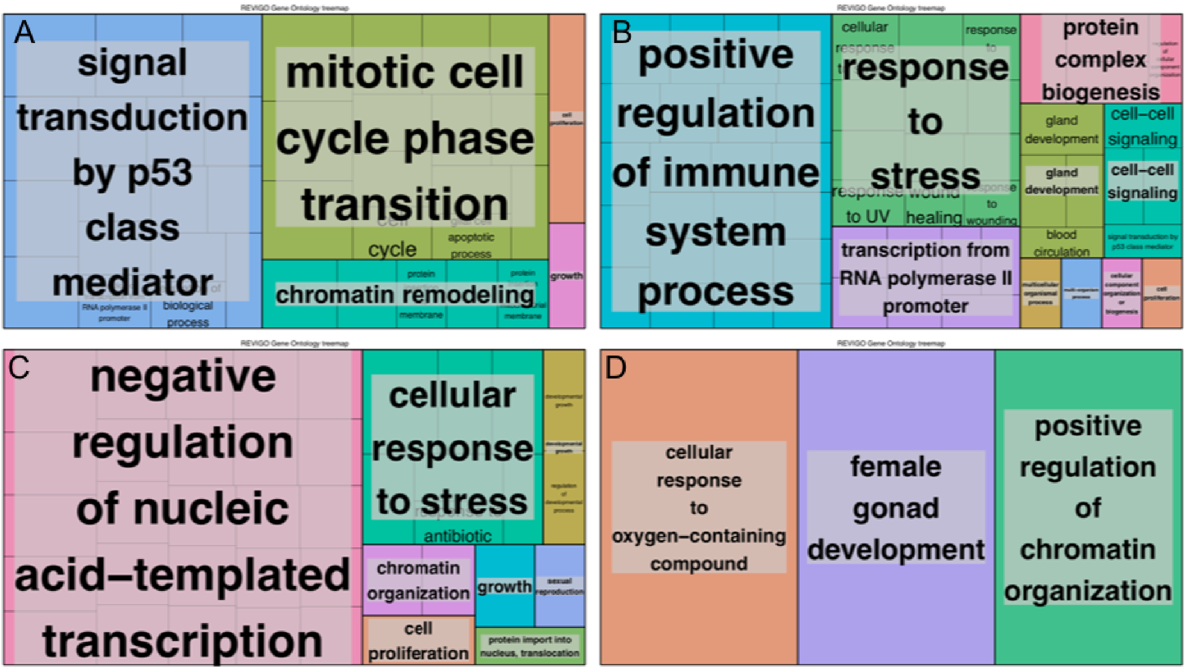


**SI Figure 7. Functional enrichment of overlapping CellAge and cancer-related genes. (A) Between CellAge senescence promoter genes and TSGs, (B) Between CellAge senescence promoter genes and oncogenes. (C) CellAge Senescence inhibitor genes and TSGs. (D) Between CellAge inhibitor genes and oncogenes.**

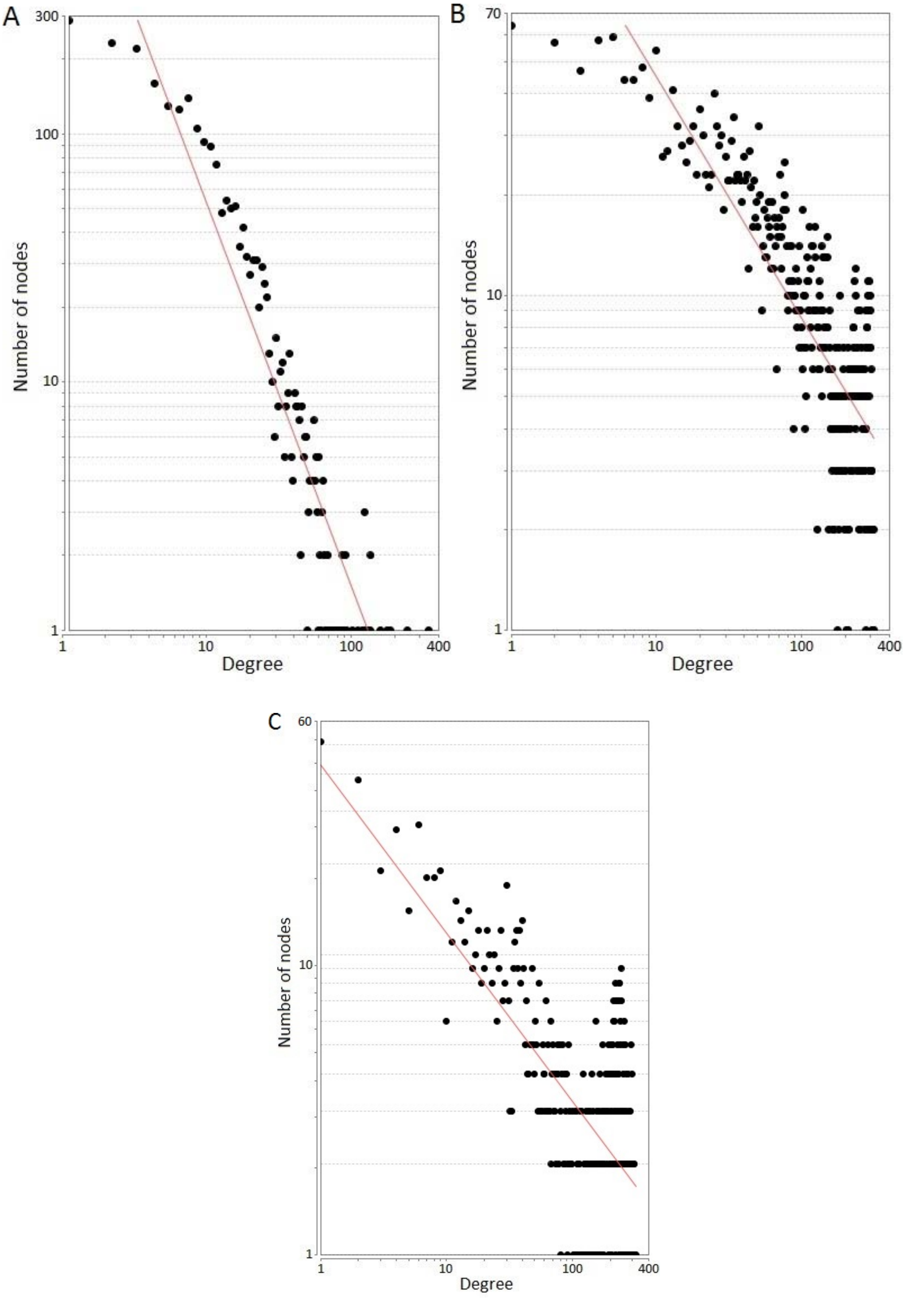


**SI Figure 8. Node degree fit to the power law distribution.** The fit of the node-degree distribution to a power law (red line) confirms the scale free structure of the networks. **(A) The Protein-Protein Interaction network** had a correlation of 0.76 and an R-squared value of 0.883. **(B) The RNA-Seq Unweighted Co-expression Network** had a correlation of 0.783 and an R-squared value of 0.630. **(C) The Microarray Unweighted Co-expression Network** had a correlation of 0.900 and an R-squared of 0.456.

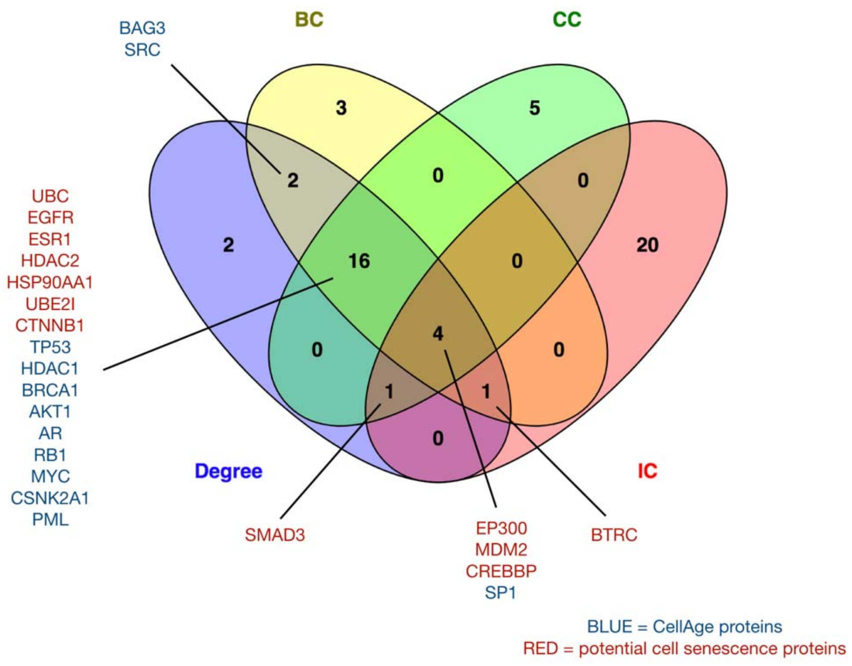


**SI Figure 9. Intersection of genes occupying multiple top positions across the network topology parameters for the PPI network.**

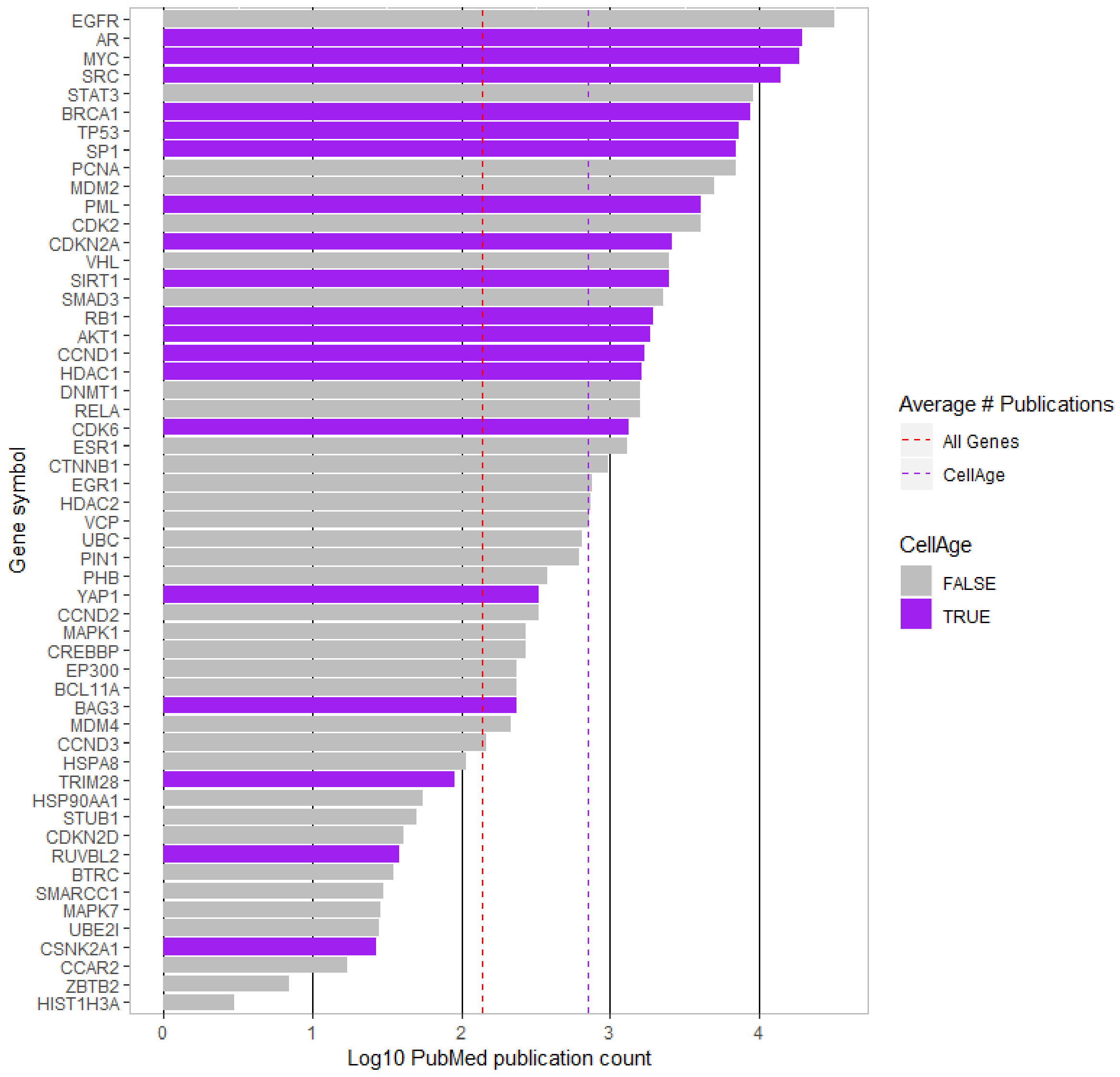


**SI Figure 10. Publication counts for high centrality genes in the PPI network.** The CellAge genes, shown in purple, generally appear in a greater number of publications. The dashed red line shows the mean number of publications for all genes, while the dashed purple line displays the mean number of publications for CellAge genes.

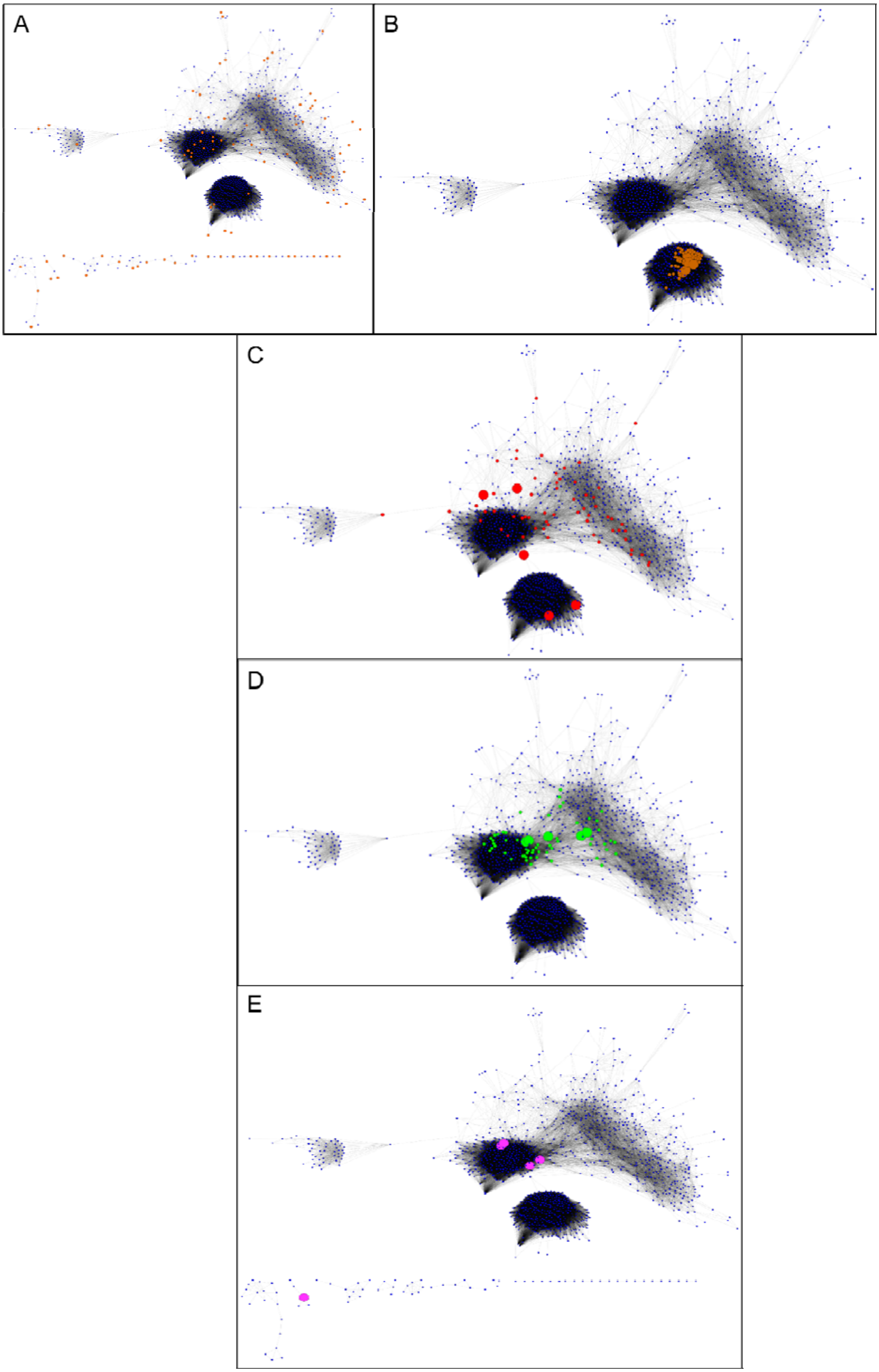
A
B
C
D
E

**SI Figure 11. The microarray co-expression network. (A) Seed Nodes of the Microarray Unweighted Co-expression Network.** The 95 seed nodes, obtained from CellAge database of senescence related genes are represented in orange, size 2X. The first interacting partners are represented in blue, size 1X. The network is divided into a main component with 1198 nodes and 18 smaller components (below) with 76 nodes in total. **(B) Top 5% Degree Microarray Unweighted Co-expression Network.** The top 5% nodes with the largest degree of the network are represented in brown, size 2X. The top 5 nodes with the largest degree (RPRML, C11orf95, NKX2-4, UTF1 and PTH2) are highlighted in brown, size 6X. The remaining nodes are shown in blue, size 1X. All the top 5% nodes were located in the main component of the network. **(C) Top 5% Betweenness Centrality (BC) Microarray Unweighted Co-expression Network.** The top 5% nodes with the largest BC of the network are represented in red, size 2X. The top 5 nodes with the largest BC (DDX54, TRIM28, SPEF1, CD164L2 and SSRP1) are highlighted in red, size 6X. The remaining nodes are shown in blue, size 1X. **(D) Top 5% Closeness Centrality (CC) Microarray Unweighted Co-expression Network.** The top 5% nodes with the largest CC of the network are represented in green, size 2X. The top 5 nodes with the largest CC (RRM1, SUPT16H, NUP205, MCM6 and DHX9) are highlighted in green, size 6X. The remaining nodes are shown in blue, size 1X. SMC4 was also at the top 5% CC. **(E) Significantly Increased Connectivity (IC) nodes in the Microarray Unweighted Co-expression Network.** The 5 nodes with significant IC with genes related to cell senescence according to CellAge database are represented in pink, size 6X. The nodes with significant IC were: HMGB2, EHD2, SMC4, LMNB1 and CKAP2. The remaining nodes are shown in blue, size 1X.

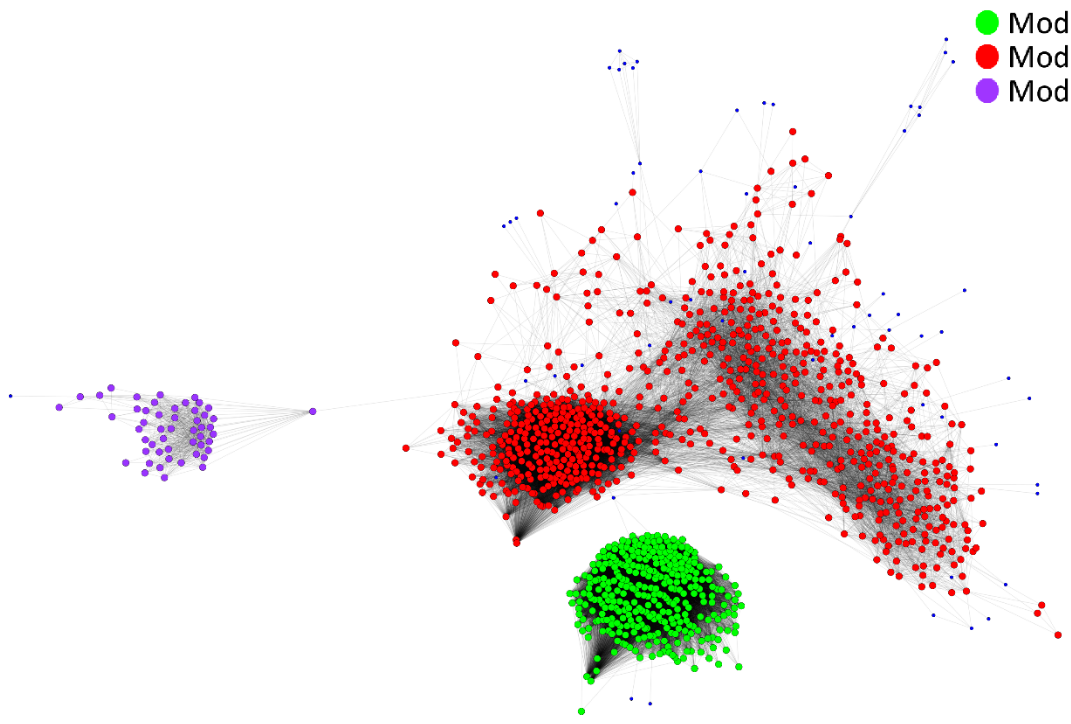

**SI Figure 12. Modular analysis of the Microarray Unweighted Co-expression Network**. On the top right, the legend links the nodes of each module to their assigned colours. There are three modules in the network. The modules are numbered in order of modularity.

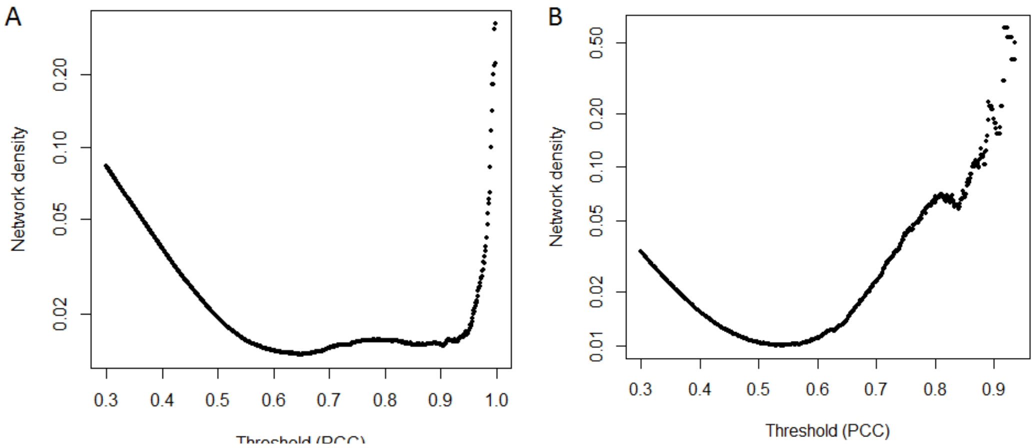


**SI Figure 13. Selection of Pearson Coefficient Correlation (PCC) threshold by network density**. The figures represent the network density of the database of interactions generated by applying each PCC threshold to **(A) the GeneFriends Database of RNAseq co-expression correlation** and **(B) the COXPRESdb Database of Microarray co-expression correlation**. The minimums of network densities are placed at 0.65 and 0.53 respectively.
